# Integrative, Segregative, and Degenerate Harmonics of the Structural Connectome

**DOI:** 10.1101/2024.02.23.581784

**Authors:** Benjamin S Sipes, Srikantan S Nagarajan, Ashish Raj

**Affiliations:** Department of Radiology and Biomedical Imaging, University of California, San Francisco, San Francisco, USA

## Abstract

Unifying integration and segregation in the brain has been a fundamental puzzle in neuroscience ever since the conception of the “binding problem.” Here, we introduce a framework that places integration and segregation within a continuum based on a fundamental property of the brain–its structural connectivity graph Laplacian harmonics and a new feature we term the gap-spectrum. This framework organizes harmonics into three regimes–integrative, segregative, and degenerate– that together account for various group-level properties. Integrative and segregative harmonics occupy the ends of the continuum, and they share properties such as reproducibility across individuals, stability to perturbation, and involve “bottom-up” sensory networks. Degenerate harmonics are in the middle of the continuum, and they are subject-specific, flexible, and involve “top-down” networks. The proposed framework accommodates inter-subject variation, sensitivity to changes, and structure-function coupling in ways that offer promising avenues for studying cognition and consciousness in the brain.

## Introduction

A central goal of systems neuroscience is to explain fundamental brain processes shared across individuals. Perhaps the most important fundamental process involves the identifying mechanisms to address the so-called “binding problem,” which asks how an integrated experience arises from segregated sensory information^[1]^. This philosophical question has been investigated in terms of brain function, where integration and segregation are ends of a continuum from synchrony to asynchrony, respectively^[2–4]^. Yet, purely functional accounts cannot explain how this integration-segregation balance emerges from the underlying biological structure. It has been proposed that white matter “structural connectivity” (SC) network topology subserves the integration-segregation balance, with integration originating from signal convergence or the “rich-club” and segregation originating from modularity^[5–7]^. However, graph metrics to measure these network properties in SC are incommensurable to a single integration-segregation continuum. Furthermore, characterizing what occurs between integration and segregation–the missing-middle–has not received sufficient attention. An integration-segregation continuum based on a fundamental network property is yet to be conceived for structural brain networks.

Understanding how integrative and segregative functional activity arises from SC depends on the “structure-function” relationship of brain networks–often described using graph theoretic and statistical measures^[8–11]^. Graph Signal Processing (GSP) extends graph theoretical approaches, providing an elegant and concrete mathematical framework to describe brain function as signal diffusion through the SC^[12,13]^. At GSP’s heart lies the graph Laplacian matrix and its decomposition into graph harmonics (eigenvectors or “gradients”^[14]^), reflecting orthogonal spatial patterns (wave functions) of a signal in the network, with each harmonic associated with an eigenvalue reflecting its graph frequency^[15,16]^. The harmonic decomposition of the graph Laplacian is tantamount to a Fourier Transform; Laplacian harmonics form the Fourier basis that describes how hypothetical brain signals would “resonate” in the structural network, thereby linking structural graph topology to functional synchrony^[12,17]^. Each harmonic is associated with an eigenvalue, and conventionally harmonics are sorted by their ascending eigenvalues.

It is therefore natural to employ the harmonic “eigenspectrum” as the organizing principle for integration and segregation, which may form the continuum we seek. This concept has recently gained currency: lower harmonics have been interpreted as integrative since they capture patterns of global synchrony, and higher harmonics have been interpreted as segregative since they capture local synchrony^[18]^. Just as low spatial frequencies in Fourier space represent broad and smooth gradients while high frequencies represent detailed, possibly localized, and noisy patterns, it has been observed that low harmonics reflect the global network structure and hence represent integrative activity while higher harmonics represent segregated network clusters^[12,19–21]^. The lowest harmonics appear to correlate with canonical functional resting-state Networks (RSNs)^[22–24]^ and were the most effective at predicting functional connectivity (FC)^[25,24]^. Accordingly, Preti et al.^[21]^ suggested dividing harmonics into two classes based on their participation in fMRI: the lowest harmonics were “coupled” to fMRI signal while all higher harmonics were “decoupled.” Sometimes these two regimes are also called “illiberal” and “liberal”, respectively^[12]^. Therefore an ascending harmonic eigenvalue ordering is a plausible means of describing the integration-segregation continuum.

However, as we show here, many aspects of the structural harmonics’ eigenspectrum make the desired integration-segregation (or “coupled-decoupled”) binary problematic and incomplete. The relationship between Laplacian eigenvalues and spatial frequency has only partial mathematical^[26]^ and empirical^[19]^ support. Higher harmonics in particular have been shown to break from strict frequency expectations^[19]^. Mathematically, graph spectra may be reasonable analogs of spatial frequencies for simpler graphs (e.g., rings and infinite or closed lattices) but not necessarily for complex networks^[26,19]^, such as the brain^[4,8]^. The notion of low harmonics being exclusively coupled to function is also imperfect: higher harmonics also significantly contribute to functional activity, such as in MEG spectra^[27,18]^ and in alignment with RSNs such as the visual network^[22]^. Thus, both theoretical and empirical evidence suggests that the coupling-decoupling dichotomy does not neatly align with the integration-segregation continuum.

Can the Laplacian harmonic spectrum serve as an integration-segregation continuum? Do the harmonics follow expectations for the integration-segregation continuum, such as reproducibility across individuals? What fundamental network property could distinguish integrative-versus-segregative harmonics? What represents the “missing-middle” of the continuum, and what is its relevance to the underlying biology and brain function?

In this study, we address these questions using a deep investigation of the structural harmonics’ eigenspectrum. First we develop a consensus SC to facilitate an ordering of harmonics across subjects. We seek the most informative ordering of the harmonics that simultaneously addresses the following criteria:

1. The ordering must place integration-like harmonics at one end and segregation-like harmonics at the other.
2. The ordering must reproduce as much as possible notions of spatial smoothness and sparsity.
3. The ordering should be reproducible across individuals in a predictable manner.
4. Harmonics placed similarly along the spectrum must share more properties with each other than those far apart.

We first show that the canonical ordering of harmonics by ascending eigenvalues only partially succeeds in fulfilling the above criteria, but that SC harmonics which deviate from classical Fourier expectations are not reproducible across subjects with this ordering. Then, we show that the above criteria can be met by aligning subject-specific harmonics to the group-level harmonics from the consensus SC (the mean SC across individuals). This new ordering reveals an underlying harmonic reproducibility related to the consensus “gap-spectrum,” a fundamental property of the eigenspectrum derived from spectral graph theory. Further, the gap-spectrum naturally partitions the harmonics into three distinct regimes: integrative harmonics have low eigenvalues but high spectral gaps; segregative harmonics have high eigenvalues and high spectral gaps; the “degenerate” harmonics have intermediate eigenvalues but low spectral gaps. The latter regime is so-named in analogy to degenerate modes observed in many physical systems, both classical^[28–31]^ and quantum^[32,33]^. This degenerate regime fills the “missing-middle,” has not previously been defined in brain science and is endowed with distinct functional relevance.

Together, the three regimes faithfully account for key group-level harmonic properties including spatial smoothness, sparsity, sensitivity to changes, similarity between individuals, and alignment to functional networks. Hence, a more complete stratification of harmonics requires a trichotomy–integrative, degenerate, segregative–instead of the popular integration/segregation or coupled/decoupled dichotomies. The proposed harmonic trichotomy aligns with prior graph theory accounts of integration-segregation while also revealing previously unreported structure-function relationships by explaining resting and task functional networks in a manner that is not possible within prior paradigms.

## Methods

### Datasets

Data in this study come from the publicly available MICA-MICs dataset of 50 healthy young adults (23 women; mean age 29.54 years)^[34]^. This dataset was generated with the micapipe connectivity pipeline, and a full description of the image processing steps can be found in their articles^[35,34]^. This dataset is not only of high quality, but it features cortical connectivity parcelled in the Schaefer atlas^[36]^ at 10 different spatial resolutions (from 100 to 1000 cortical nodes), with each parcellation including an additional 6 bilateral subcortical regions and the bilateral hippocampus (14 additional nodes total). We used networks weighted by classical streamline count connectivity (hereafter “SC”) as well as the time series from resting-state functional magnetic resonance imaging (fMRI) scans (TR = 600ms, 695 time points). Most analyses were performed in MATLAB while the Neurosynth analysis was performed in Python.

To ensure that our main findings were replicable, we used another dataset of SC collected with 220 healthy subjects (ages 18-75; mean(std) = 39.1 *±* 16.8; 119 women). This replication dataset was previously used as the healthy control sample for a recent study analyzing differences in tinnitus^[37]^. Subjects in this replication dataset had more diverse age demographics, were imaged with different scanning parameters, were processed with a different connectivity pipeline, and were parceled in a different atlas, the Brainnetome atlas^[38]^ (Supplement Section 1).

### Connectome Harmonics

For each subject’s SC, we first normalized their SC matrix to a unit Frobenius norm, then we computed the degree-normalized Laplacian:

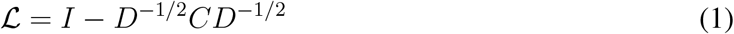

where *I* is the identity matrix, *D* is a diagonal degree matrix of the SC matrix (*C*). We then computed the eigen-decomposition of the resulting Laplacian matrix.

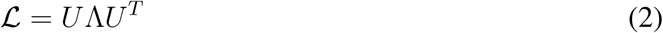

In (2), *U* is the orthonormal matrix of harmonics in each column and Λ is the diagonal matrix of eigenvalues associated with each harmonic. Additionally, we will henceforth use *N* to denote the number of regions for the SC parcellation.

### Harmonic Frequency Analysis

Determining the precise spatial frequency of each harmonic is not straightforward since the brain’s harmonics live in the brain’s 3-dimensional space, and a 3D-FFT does not have an obvious single feature representative of the full volume’s frequency content. We instead used four measures that relate to spatial frequency and analyzed their relationship to eigenvalues. For the below measures we first apply a modest threshold (*T* = *±*0.001) to eliminate spurious low-amplitude fluctuations around zero.

Harmonic sparsity is measured as the fraction of regions where the harmonic’s amplitude is zero. In classical Fourier analysis, the sparsity of all harmonics in a Fourier transform are zero (i.e., all frequencies have infinite support). Therefore, SC harmonics with high sparsity defy Fourier-based intuitions for harmonics.

We used MATLAB’s `zerocrossrate` function to measure the rate at which the SC harmonic changes polarity (crosses zero), with high frequencies defined by a high rate of polarity switching. This measure assumes an ordering of regions such that adjacent regions are also adjacent across the harmonic’s indices. This assumption is generally appropriate, as is clear in the Regional Adjacency matrix that regions close in space are generally adjacent in the regional ordering (Supplementary Figure S3).

Harmonic Network Zero Crossings as a measure of harmonic frequency was proposed by Huang et al.^[19]^, which accounts for Laplacian harmonics being graph cuts on the network:

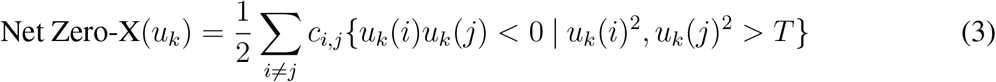

Descriptively, the network edges between regions with opposite signs in the harmonic are summed together, which is the graph cut weight described by the harmonic. This measure can be considered a proxy for the “frequency” of the harmonic with respect to the underlying network since higher frequencies will have less optimal (highly weighted) graph cuts. However, one known difficulty to this approach is that the areas of low harmonic amplitude fluctuate spuriously around zero, and therefore the above threshold (*T*) is useful to show graph cuts between regions that meaningfully participate in the harmonic. To see results across thresholds (Supplementary Figure S4).

We propose a “Spatial Roughness” measure for a given harmonic’s frequency that uses a regional adjacency matrix (*A*), where the entry *a*_*i,j*_ is a reciprocal function of the distance between adjacent voxels in MNI space between regions *i* and *j*. Specifically,

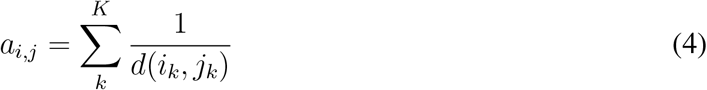

In this equation, the summation is over all *K* voxels on the boundary between regions *i* and *j*, and *d*(*i*_*k*_, *j*_*k*_) is the euclidean distance between the adjacent voxels *i*_*k*_ and *j*_*k*_.

Our formulation for Roughness then becomes:

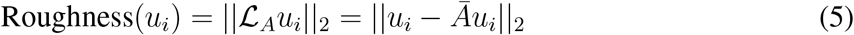

where *ℒ*_*A*_ is the degree-normalized Laplacian of the regional adjacency matrix (*A*), *u*_*i*_ is the *i*^th^ eigenvector of the SC Laplacian, Ā is the degree-normalized adjacency matrix 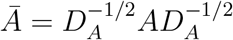, and (||*·*||_2_) is the euclidean (L2) norm of the result. The term ||*u*_*i*_ − *Āu*_*i*_||_2_ captures roughness by measuring how much the harmonic signal *u*_*i*_ deviates from its weighted average based on the regional adjacency, effectively quantifying local signal variability. Since *A* is defined only within each hemisphere, we compute the Roughness separately for each hemisphere then average between them. Spatial roughness is high when adjacent regions in the brain are dissimilar in both their harmonic’s signal amplitude and polarity (i.e., sign).

### Harmonic Matching

We computed eigenvectors for each subject’s SC individually as well as for a consensus SC, which was formed by taking the average across all subject’s networks, computing the normalized Laplacian (1), then obtaining its eigen-decomposition (2). We then matched each subject’s eigenvectors to those of the consensus network with a bipartite matching max-flow algorithm^[39]^ implemented in MATLAB (the `matchpairs` function). Mathematically, this function finds a permutation that maximizes the trace of |*U*^*T*^ *V* |, where *U* is the subject’s eigenvectors, and *V* are the consensus eigenvectors. Let us define a permutation vector *p* such that it maximizes:

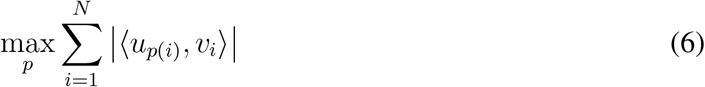

Where |*·*| denotes an absolute value, and *u*_*p*(*i*)_ is the column in *U* corresponding to the *i*^*th*^ index in the permutation *p*. By matching individual subjects to the consensus SC, we retain an ordering of ascending eigenvalues with respect to the consensus SC.

Note that others have recently performed eigenvector alignment using Procrustes^[40]^; however, we refrained from this approach to preserve as much of the subject-specific features in the harmonics as possible, since understanding the subtle harmonic variation across subjects is a key question evaluated in this work.

### Inter-subject Agreement

To evaluate inter-subject agreement between both unmatched and matched harmonics, we computed all possible combinations of subject-specific consensus-matched harmonic dyads (subject_*i*_, subject_*j*_) and took the absolute value of each diagonal:

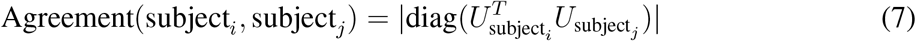

Agreement between subjects *i* and *j* is a vector in ℝ^*N*^, and we computed agreement for all pairs of subjects in our dataset. Note that since these harmonics are orthonormal, subjects that share identical harmonics will have an Agreement equal to 1 for those harmonics, and fully orthogonal harmonics will have an Agreement equal to 0.

Separately from inter-subject agreement, we analyzed inter-subject variation within each region across harmonics. After matching to the consensus SC harmonics, we re-scaled subject-specific absolute-valued harmonics so that each was on the closed interval [0, 1], then we took the variance across all subjects followed by the mean across all harmonics.

We replicate our primary inter-subject agreement analysis in two complementary ways. First, we replicate across parcellation resolutions with the Schaefer atlas (114-1014 parcels) in the same MICA-MICs dataset (Figure S5). Second, we replicate in an entirely separate dataset with 220 subjects processed with different MRI acquisition parameters, a different SC pipeline, and parcelled in the Brainnetome atlas with 246 regions (210 cortex plus 36 subcortex)^[38]^ (Supplement Section 1). This held-out dataset has been previously published here^[37]^.

### Eigenvalue Gap-Spectrum

In spectral graph theory, harmonic eigenvalues index different network topologies^[41]^, and the difference (or “gap”) between ascending eigenvalues can have specific meanings–for example, the gap between the smallest and second smallest eigenvalues quantifies the network’s overall connectivity^[15,42,43]^. In general, each harmonic captures dimensions of the networks structure, and the closer two eigenvalues are to each other (i.e., the smaller their gap, the more they are degenerate), the closer the associated harmonics are to describing the same type of network configuration, i.e. network motifs^[41]^ and symmetries^[44]^. More generally, the “gap” between any two eigenvalues is:

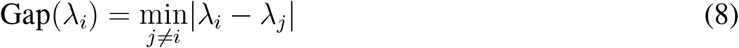

Spectral gaps indicate the “degeneracy” between two harmonics, where harmonics with the same eigenvalue (i.e., a gap equal to zero) are considered fully degenerate. Degenerate eigenvectors have a sense of “flexibility” since they represent symmetries in the network^[44]^, they are related to network synchonizability^[45]^, and they can form linear combinations to produce new degenerate eigenvectors. Here, we analyzed the SC harmonic eigenspectrum through what we’ve termed its “gap-spectrum,” which is analogous to the derivative of the ascending eigenvalues with respect to their index, and which represents a measure of harmonic degeneracy. However, because derivatives are known to amplify noise, to obtain a smooth gap-spectrum for all harmonics, we first fit an order 10 spline with 3 knots to the SC eigenvalues and then computed the first analytical spline derivative as the gap-spectrum. We hypothesized that the first-order gap-spectrum (measuring harmonic degeneracy) would be related to various properties of harmonics.

### Defining Harmonic Regimes

We sought to define regimes of harmonics to encompass empirical group-level features across subsets of harmonics. Ideal regimes should group together integrative, degenerate, and segregative harmonics in a way that retains relationships with harmonic spatial frequency and inter-subject agreement. Importantly, while past work has classified harmonics based on their participation in empirical functional activity^[21]^, we sought a classification feature agnostic to brain function and instead based on a natural property of structural connectivity: the normalized Laplacian eigenspectrum.

Since the gap-spectrum accounts for harmonic degeneracy, it becomes a natural way to classify harmonics into three regimes: those with low eigenvalues and high gap-spectrum, those with medium eigenvalues and low gap-spectrum, and those with high eigenvalues and high gap-spectrum. The simplest approach is to set a threshold on the gap-spectrum, however this choice would be somewhat arbitrary. In an attempt to impart some rigor and reproducibility, we instead sought the natural points of inflection in the gap spectrum. Specifically, we set the lower and higher bounds of the degenerate regime as the first local maximum and last local minimum in the gap-spectrum derivative (Supplementary Figure S5).

While it may be sufficient to classify harmonics based solely on the consensus SC eigenspectrum, we sought to ensure the classification was appropriate for all subjects by first classifying each harmonic based on the subject’s individual gap-spectrum, then we matched those subject-specific harmonics to those from the consensus SC (Equation 6). Therefore, for each harmonic, there was a “vote” from each subject for that harmonic’s regime classification. We defined the final “consensus regimes” based on the majority vote. We used this regime assignment for all subjects in subsequent analyses.

### Stability of Harmonics

We hypothesized that the stability of a harmonic to network changes would be related to the gap-spectrum. We perturbed each subject’s natural SC matrix by adding a matrix of Rician noise *R*(*σ, ν*)^[46]^ with *ν*= 0, which reduces to a Rayleigh distribution^[47]^, and with a scale parameter *σ*, which is related to the distribution’s standard deviation. We chose this distribution given its relevance to MRI imaging^[46,47]^. We then define our perturbed connectivity matrix as:

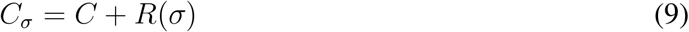

We then computed the Laplacian harmonics for each *C*_*σ*_ (i.e.,*U*_*σ*_), matched *U*_*σ*_ to the same subject’s unperturbed harmonics (*U*) using Equation 6, and then computed their perturbation as:

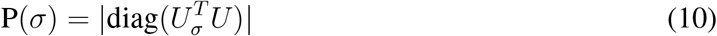

Thus, for each *σ* we obtained a measure of similarity for every harmonic across subjects. Drawing inspiration from perturbation theory and sensitivity analysis^[48,49]^, we can find the stability of harmonics as:

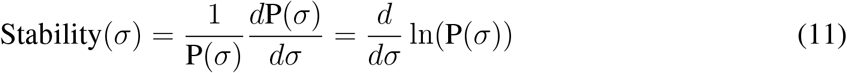

We evaluated this stability numerically in the range *σ* = [0, 0.002] and step size *dσ* = 2.5 *×* 10^−4^ and finding the slope between *dσ* and ln(*P* (*σ*)). We computed stability separately for each harmonic and then compared across subjects to the gap-spectrum.

### Harmonic Regime SC

Given our interest in how different SC features are reflected across harmonics, we sought to analyze the graph theoretical properties of a low-rank network reconstruction of SC from a limited subset of harmonics. We hypothesized that different subsets of harmonics would embody different graph theoretical properties conventionally related to integration and segregation.

To accomplish this, consider the spectral decomposition of the graph Laplacian (2). To reconstitute a low-rank structural connectivity matrix, we first compute the following truncated spectral decomposition:

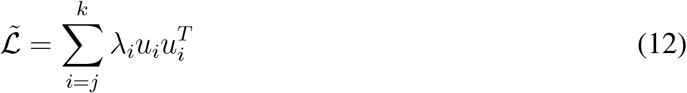

Where the interval [*j, k*] is a subset of harmonics. Then reconstituting the underlying low-rank SC becomes:

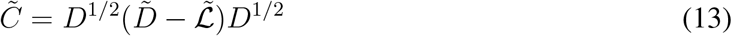

Where here *D* is the original diagonal degree matrix of *C*, and 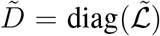. However, since the spectral summation is truncated, there will be negative weights in 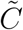 reflecting the connections that were excluded from the truncation; therefore, we threshold all 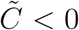 to be zero. We also provide a more detailed derivation of 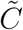 in the Supplement Section 5.

To analyze the integrative and segregative properties of 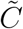 for different subsets of harmonics, we used functions implemented in the Brain Connectivity Toolbox^[50]^ to compute the following graph theoretic measures for weighted undirected networks: efficiency, small worldness, modularity, and coreness. We use the small worldness measure^[51]^, which is defined as:

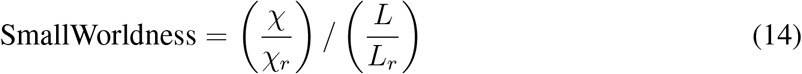

In this equation, χ is the network’s clustering coefficient, *L* is the network’s characteristic path length, and χ_*r*_ and *L*_*r*_ are the clustering coefficient and characteristic path length for a randomized network, respectively. To produce randomized networks, we used the Brain Connectivity Toolbox (<u>null model und sign</u>). Efficiency and small worldness capture more integrative network topologies while modularity and coreness capture segregative network topologies^[7]^. To analyze these network properties across regimes, we normalize each 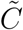 to give it a unit Frobenius norm, 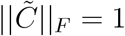.

### SC Harmonics’ Participation in Function

To investigate how harmonics are related to the brain’s function, we define harmonic participation by how known functional resting-state networks, meta-analytic task networks, and empirical resting-state time series align with each harmonic’s power, defined as:

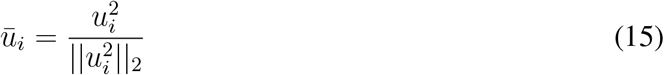

Here, the ||*·*||_2_ denotes the L2-norm. We then concatenate these vectors into the columns of a matrix 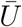.

We use 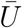 to analyze the spatial power distributions of harmonics (in analogy to the”Born rule”^[52]^), and we analyze this “harmonic power” relationship to brain function. In other words, we ignore harmonic polarity (i.e., sign) and instead analyze how the harmonic’s power relates to 1) resting-state networks, 2) meta-analytic task networks, and 3) empirical resting-state time series. We describe each of these three functional analyses below.

Alignment with Resting-State Networks: Using the Schaefer atlas, each node in the network was labeled according to the 7 canonical resting-state networks^[23]^, with an additional label for the 14 subcortical regions (which we’ll consider analogously to the other RSNs for this analysis). To evaluate harmonic participation with RSNs, we first created a binary matrix *Q* with rows equal to the number of network nodes and eight columns (one row for each RSN). For each entry in *Q, q*_*i,j*_ = 1 if node *i* belongs to RSN *j*, and *q*_*i,j*_ = 0 otherwise. We then computed RSN participation matrix for each subject’s matched harmonics 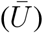 as

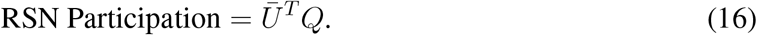

We are most interested in the relative participation between various RSNs, so we row-normalize the RSN Participation matrix by the maximum value in each RSN. For the participation without row-normalization, see Supplementary Figure S6.

Alignment with Meta-analytic Task Networks: To evaluate the correlation between SC harmonics with meta-analysis task activation maps, we exported the power harmonics 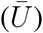 from the 214 node network for each subject into NIFTI files in MNI space. We then used the NiMARE toolbox in Python to compute the correlation between the eigenvectors and Neurosynth task activation maps for 146 Latent Dirichlet Allocation (LDA) topics drawn from the LDA-400 set (Neurosynth version-7). We included topics related to cognitive processes while we excluded topics related to methodology or clinical conditions. We then averaged subsets of the 146 topic terms to obtain 24 separate cognitive domains similar to those applied in previous meta-analysis studies using Neurosynth^[14,21]^. For a full list of LDA terms and their cognitive domain assignments, see Supplementary Data.

Empirical Resting-State fMRI: Finally, we analyzed the properties of harmonic power vectors in empirical resting-state fMRI time series. This projection follows a similar procedure as previous works^[53,21,24,54]^, but we instead use the power harmonics (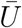, Equation 15). Given a time series (*s*(*t*) ∈ ℝ^*N*^), we define the projection:

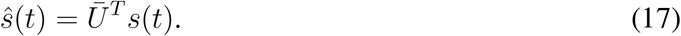

We refer to this as a projection “onto the harmonic power vectors.” Note that if we used *U* instead of 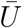 this equation would be a graph Fourier transform. To study the functional properties of the projected signals, we analyzed the signal’s energy, temporal diversity (sample entropy), spatial entropy (harmonic diversity), and temporal dynamics.

The energy of a signal *(ŝ*_*i*_(*t*)) is defined as the L2-norm squared across time, which by Parseval’s theorem is equivalent to the integral of the signal’s spectral density^[55]^. We computed the energy for each time series projection signal (*ŝ*_*i*_) to compare across harmonic regimes. We further assessed the regional distribution of harmonic energy by projecting the energy back into the region space.

We used a measure of temporal entropy, known as Sample Entropy (SampEn,^[56,57]^), which measures the self-similarity of signal sub-sequences across time. Intuitively, this measures the signal’s temporal diversity. We used the MATLAB function sampen to compute the signal’s sample entropy^[58]^. Sample entropy requires defining a scale parameter *m* and a similarity tolerance parameter *r*. For biological signals, it has been suggested that *r* = 0.2^[57]^, which we have followed here. There is no standard for choosing *m*, but larger *m* accounts for more dynamics in the data, and it is recommended to be chosen to match the dynamic timescale of interest in the signal. Accordingly, we chose *m* = 5 which corresponds to 5 TRs (i.e., 3-seconds) in the signal. Each harmonic time series *ŝ*_*i*_ has one value of sample entropy, and we averaged sample entropy across regimes to evaluate whether regimes differ in their temporal diversity.

Following the measure of repertoire diversity introduced in^[54]^, we defined a similar measure we call “harmonic diversity” for the projected time series:

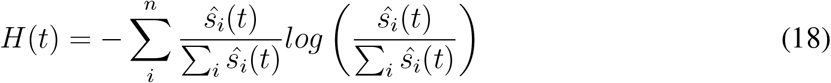

Since this measure is influenced by the number of harmonics in a given regime, we normalized by the maximum possible diversity for each regime *log*(*n*), where *n* is the number of harmonics in the regime. We compute this measure of harmonic diversity within each regime, then average across time to compare across regimes.

Finally, we analyzed the dynamics of harmonics by analyzing the relative signal strength of each harmonic regime, which can be visualized in a ternary plot. We analyzed these dynamics by the mean relative contribution of each regime as well as by analyzing the first principle component to show the main axis of dynamic fluctuation.

Ternary Plot Analysis: Since we used the gap-spectrum to identify three regimes of harmonics, we interpreted some of the above analyses using ternary plots. Ternary plots are a visualization technique to show the relative contributions of three different phenomena in a mixture^[59]^. We generate ternary plots in MATLAB using the code provided here^[60]^. As such, the values displayed in a ternary plot are relative “quantities,” and therefore each point represents a coordinate along three dimensions that sum to 1. In our work, the quantity is the amount of participation in each of our three harmonic regimes. Importantly, the ticks on the axes of a ternary plot show the direction of the axes through the plot. We have included labels for each axis in our plots to indicate which axis corresponds to which harmonic regime, and we color data inside the ternary plot according to how close each datum is to a mono-regime vertex. To limit the influence of spurious correlations in this analysis, we applied an 80^th^ percentile thresholded (across all subjects) to both the RSN participation strength and Neurosynth correlations, we then averaged across subjects to obtain group-level harmonic participation maps for the RSNs and Neurosynth.

### Statistics and Reproducibility

In this study, we used three types of statistical tests: Pearson’s correlation, Analysis of Variance (ANOVA), and two-sided t-tests. For comparing harmonic properties with the gap-spectrum, we used a Pearson’s correlation and show the distribution in scatter plots with trend lines. When we compared the gap-spectrum with harmonic properties in Figure 3B, we correlated the median value from across 50-subjects to the gap-spectrum, with the number of data-points in the correlation equal to the number of harmonics in the regime. All R-values for correlations are shown as figure insets. When we compared the stability of harmonics to the gap spectrum, we computed the Pearson’s correlation for each subject’s individual stability values compared to the gap-spectrum, which has a sample equal to the number of harmonics (n=214). In Figure 4C, we show the distribution of R-values from that analysis as an inset, and a single scatter plot aggregating all values together as the main visualization. In the analyses where we compared across regimes, we first conducted a 1-way ANOVA to ensure group-level differences, then we used two-sided t-tests to determine the between-group significance. These analyses compared the full sample of 50 subjects represented by the indicated measures from different regimes. Exact values for reported statistics is contained in the Supplementary Data 2. In the Supplement Section 7, we show some additional statistical tests that verify the significance of the matching, in which we show the significance level as −*log*(*p*) as well as the Bonferroni-corrected significance level as a red line in the Manhattan plot. The original data and MATLAB code is provided for all primary analyses (https://github.com/RajLab-UCSF/IntDegSeg), ensuring the complete reproducibility of our results.

## Results

Here, we present results that investigate whether SC harmonics may constitute an integration-segregation continuum in the brain. We begin by showing that harmonics deviate from strictly ascending frequencies and have an integration-to-segregation like structure. We then show that integrative and segregative harmonics are shared across individuals, which would indeed be expected for any candidate brain mechanism to resolve the binding problem. Then, using the gap-spectrum, we define integrative, segregative, and degenerate regimes precisely, and we further assess regime relationships to graph frequency and graph theoretical metrics. We conclude by investigating the functional relevance of the three regimes in multiple ways, including stability to changes, alignment with resting-state networks, task networks, and empirical resting-state time series.

### SC harmonics’ spatial frequencies deviate from Graph Fourier intuitions

SC normalized Laplacian eigenvalues are similar across subjects and are close to those from the consensus SC (Figure 1A). We also show that the lowest harmonics in the consensus SC qualitatively matched with those reported in prior works^[61,54,40]^ and that higher harmonics are increasingly localized (Figure 1B). Quantitatively, we investigated properties of SC harmonics related to classical Fourier intuitions, including sparsity and three spatial frequency measures: zero-cross rate, network zero-crossings^[19]^, and roughness. First, we show that the harmonics with the highest eigenvalues are sparse (Figure 1C). Next, based on Fourier intuitions, we may expect a monotonic relationship between harmonics’ eigenvalues and spatial frequency metrics, yet all three measures show non-monotonic relationships, with the greatest deviation for the highest eigenvalue harmonics (approximately when *λ* ≥ 1; Figure 1C).

**Fig 1:**
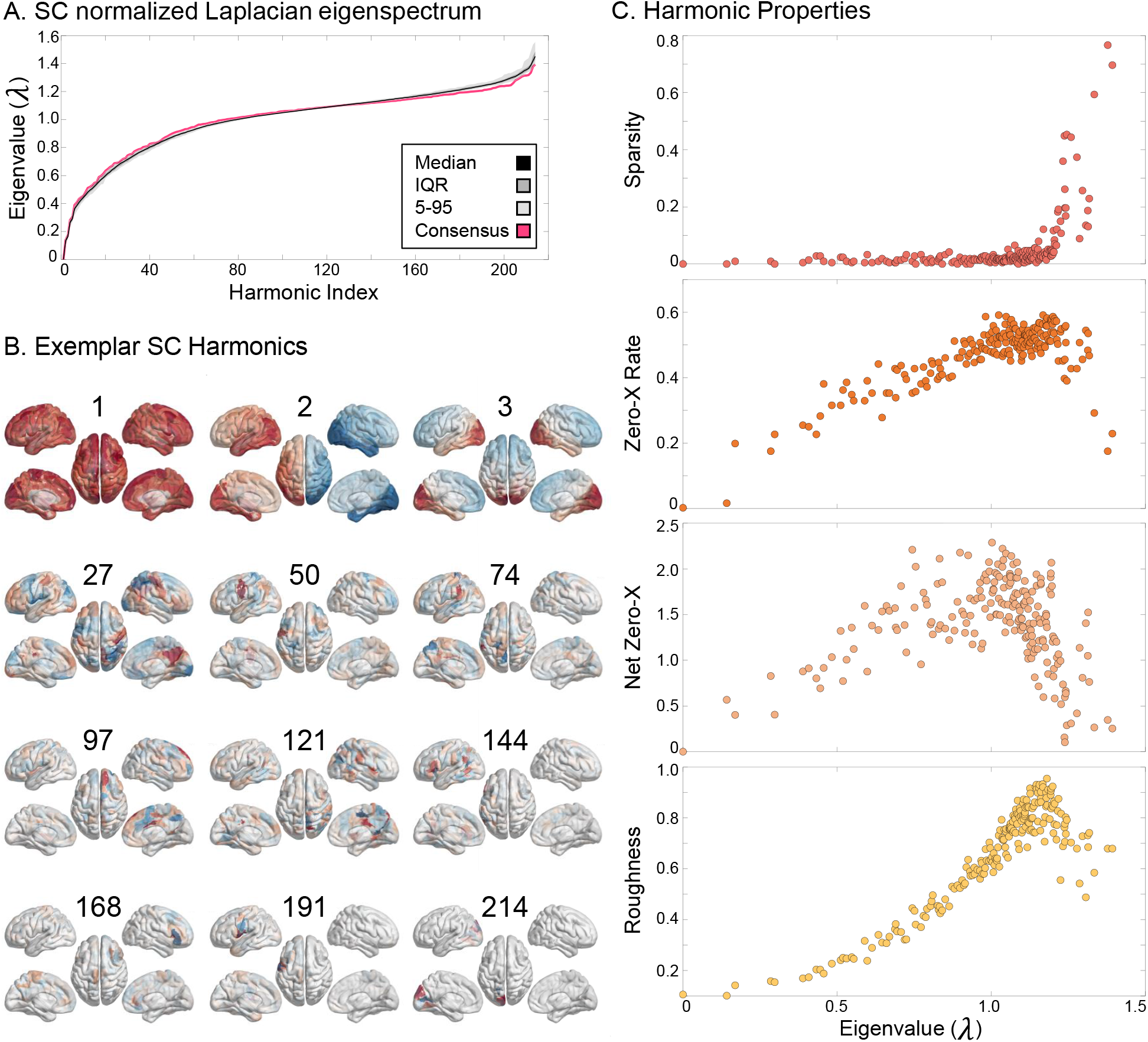
SC Harmonic Spectrum. (A) We show the normalized Laplacian eigenvalue spectrum for the SC network with 214 regions. In rose is the eigenvalue spectrum from the consensus (group-averaged) SC, and overlain on top in black is the median eigenvalue spectrum across all 50 subjects with shaded group percentiles. (B) We show several harmonics from the consensus SC with their harmonic index labeled above each plot. Notice that the harmonics become sparse and localized as eigenvalues increase. (C) We show four properties of the consensus SC harmonics: (top to bottom) sparsity, zero crossing rate, network zero crossings, and spatial roughness. The highest eigenvalue harmonics show spatial frequency characteristics that are non-monotonically related to their eigenvalues.

These results also replicated in our independent dataset with different MRI acquisition parameters and used a different SC generation pipeline (see Supplement Section 1, Figure S1). For completeness, we also analyzed the temporal frequency characteristics of a resting-state time series projections (Supplementary Figure S8), finding that higher harmonics are not necessarily related to higher temporal frequency in fMRI.

Taken together, we find only partial support that SC Laplacian eigenvalues correspond to spatial frequency. While there is undoubtedly a relationship, it is not monotonic. Instead, these results suggest the presence of global “integrative” harmonics with low eigenvalues and sparse “segregative” harmonics with high eigenvalues. Additionally, this non-monotonic relationship implies that sorting all subjects harmonics by their ascending eigenvalues could result in misalignment between subjects.

### Integrative and segregative harmonics are similar across subjects

Next, we used a measure of inter-subject agreement (7) to test whether subjects had similar harmonics for both whole-brain (integrative) and sparse (segregative) ends of the eigenspectrum. When all subjects’ harmonics are sorted by their ascending eigenvalues, we only observe high agreement for the first few harmonics (Figure 2 A & B). However, matching harmonics to the consensus SC revealed that many subjects indeed shared harmonics at both ends of the consensus eigenspectrum (Figure 2 C & D).

**Fig 2:**
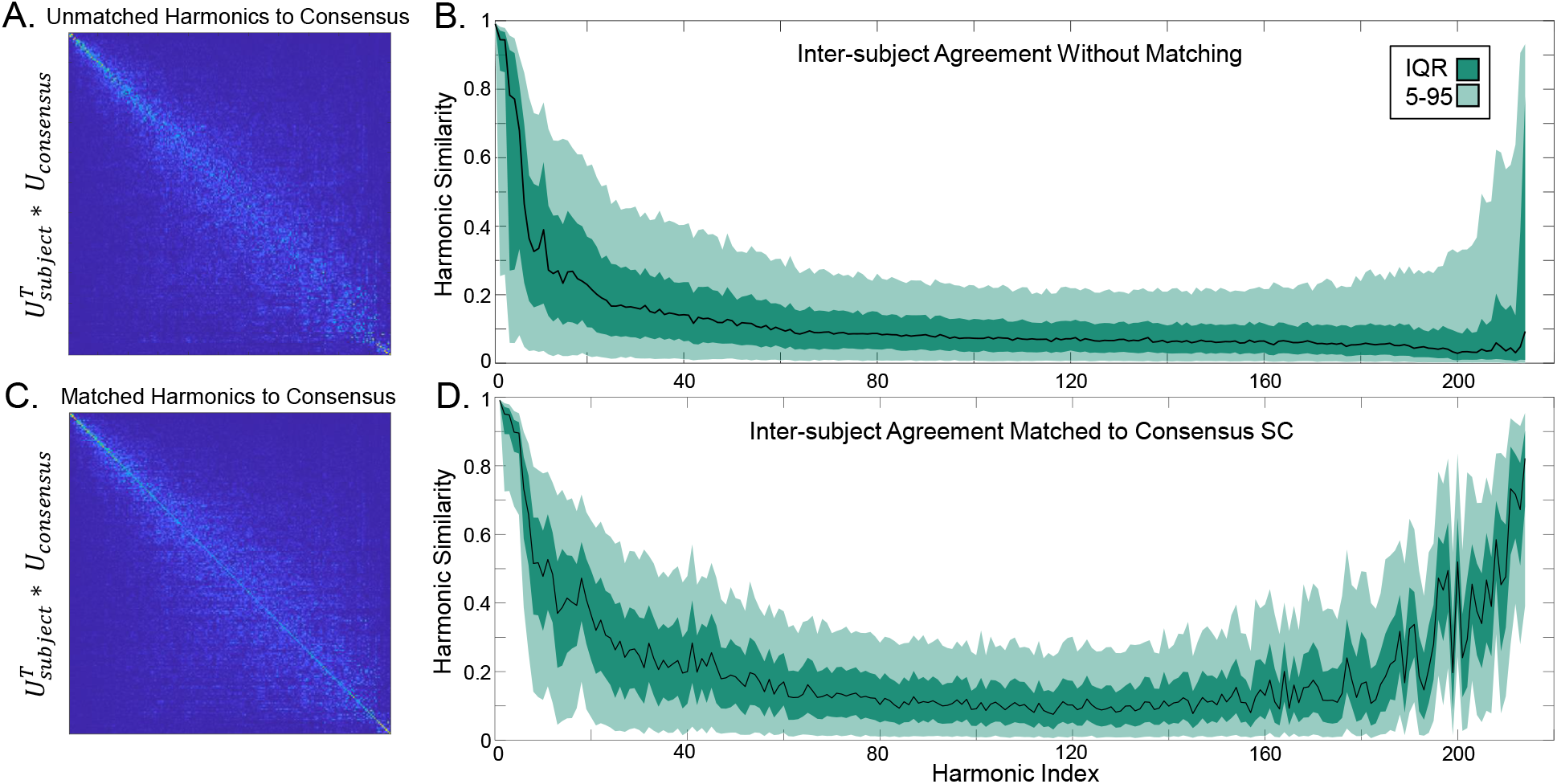
Harmonic Inter-subject Agreement. (A) Harmonics do not show a natural alignment to the consensus SC, and (B) inter-subject agreement, measured as the harmonic similarity across all combinations of subjects, showed that most subjects are misaligned for most harmonics. But (C) matching harmonics can maximize this similarity on the diagonal, and (D) following harmonic matching, we observe that subjects share many of the same integrative and segregative harmonics. In B & C, the black line indicates the median, while the shaded regions indicate the interquartile range (IQR; darker) and 5th to 95th percentiles (lighter).

We found this result to be consistent across all Schaefer parcellation scales (Supplementary Figure S9). This result also replicated in an independent dataset with different MRI acquisition parameters, a different SC generation pipeline, and a different brain atlas (see Supplementary Figure S2). Furthermore, we evaluated the Bonferroni corrected significance between the two agreement distributions and found that nearly all harmonics showed highly significant agreement improvement after matching (Supplementary Figure S6). We then generated null-model networks with randomized edge weights from empirical SC networks and found that these randomized networks were unable to match. Empirical matching showed highly significant agreement compared to randomized harmonics (Supplementary Figure S7).

We also sought to evaluate which regions had the most inter-subject variability across harmonics, finding that sensory cortices and the orbitofrontal cortex had the least inter-subject variation while trans-modal regions had the most inter-subject variation (Supplementary Figure S10). See the Supplement Sections 11 and 12 for other analyses on permutation length and distance (Figure S11) and quality of consensus matching (Figure S12).

### Eigenvalue gap-spectrum as a theoretical explanation for harmonic properties

The harmonic inter-subject similarity analysis suggests there are at least three kinds of harmonics: those with low eigenvalues and high inter-subject agreement, those with high eigenvalues and high inter-subject agreement, and those with eigenvalues near *λ* = 1 and low inter-subject agreement.

We hypothesized that change-points in the “gap-spectrum” curvature (see Methods) would naturally partition harmonics into regimes that follow the empirical relationship with inter-subject agreement while preserving relationships with harmonic frequency. We first found regime transition points for each subject individually, then we matched subject-level harmonics to the consensus SC, and finally we determined a consensus regime assignment (Figure 3A middle). We found the lowest regime to span harmonic 1 to 26, the central regime to span harmonic 27 to 187, and the highest regime to span 188 to 214.

**Fig 3:**
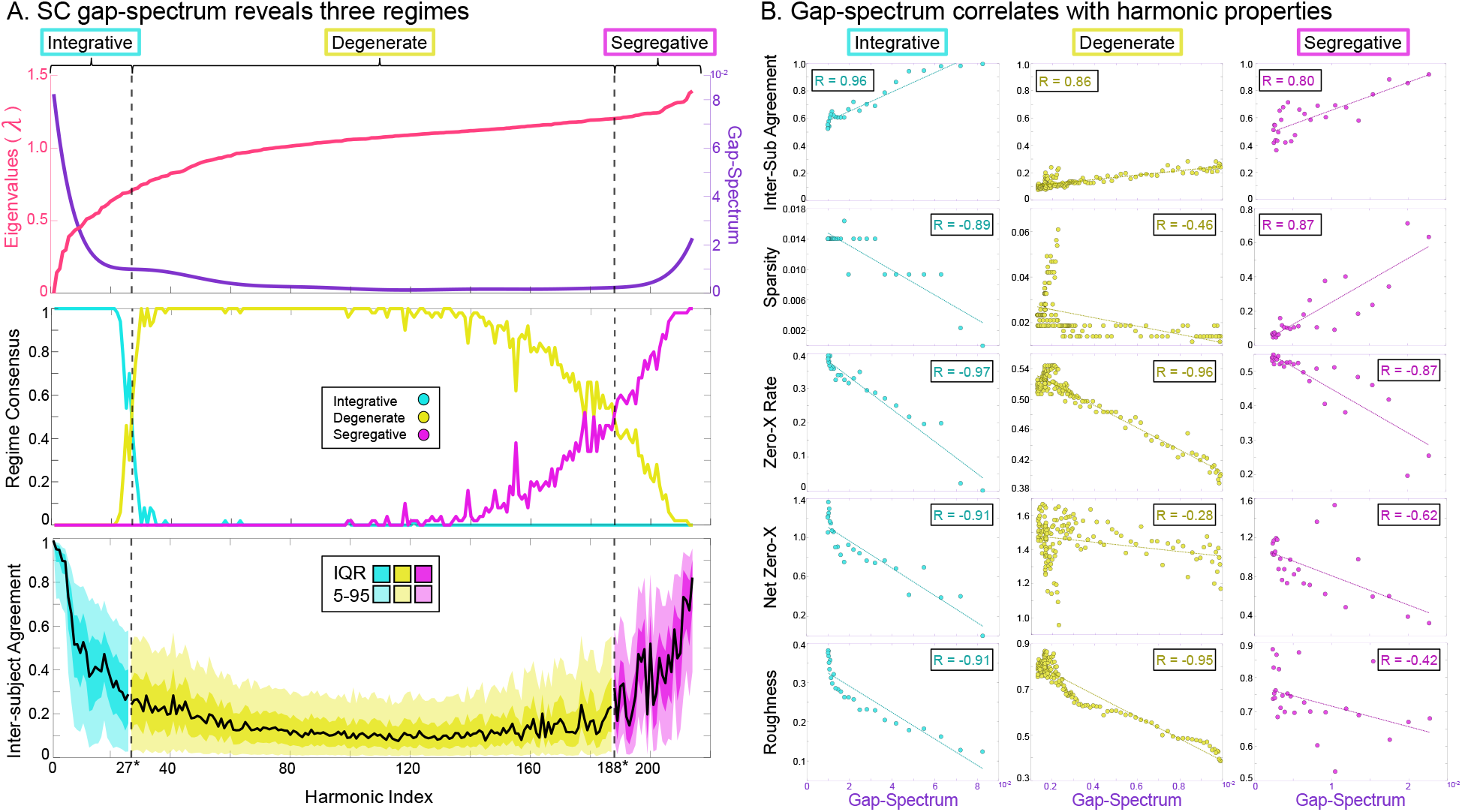
Defining Three Harmonic Regimes. (A) We show the consensus SC Laplacian eigenspectrum in rose and the analytical gap-spectrum in violet. Below this plot, we show the fraction of subjects who agree on each harmonic regime assignment. We define the final regime boundaries based on the majority crossing point. The lowest panel shows inter-subject agreement (same as shown in Figure 2D) colored by the three regimes. (B) We show the population median values for the four harmonic frequency measures compared to the gap-spectrum. All measures show strong and significant correlations within the identified regimes and the gap-spectrum.

Our results support the hypothesis that the gap-spectrum across subjects can indeed partition harmonic inter-subject agreement in an intuitive fashion while preserving relationships to grouplevel harmonic frequency (Figure 3B; all *p <* 0.05). Based on our harmonic frequency analysis, we term the lowest regime as “integrative” since these harmonics show low-frequency wholebrain spatial configurations, and we term the highest regime “segregative” since these harmonics form segregated clusters. Now, based on the gap-spectrum analysis, we term the central regime “degenerate” to reflect that these harmonics have small spectral gaps characteristic to degenerate modes of a system.

### Stability of harmonics relates to the gap-spectrum

We next sought to evaluate how SC harmonics change in response to small network changes, expecting that different regimes would show different responses to perturbation. Degenerate harmonics with eigenvalues close to *λ* = 1 represent recurring network configurations (or “motifs”)^[41]^, and they have mathematical “flexibility” since linear combinations of degenerate harmonics can produce another degenerate harmonic in the same eigen-subspace. Based on this theory, we hypothesized that the stability of harmonics would be related to the gap-spectrum and that degenerate harmonics would be the least stable since small perturbations could easily change the network’s motif structure.

We indeed found that degenerate harmonics were the most sensitive to perturbation while integrative and segregative harmonics were more stable (Figure 4A). As perturbation *σ* increased far beyond the mean network weight, we found that all regimes approximated an exponential decay (Figure 4A inset).

**Fig 4:**
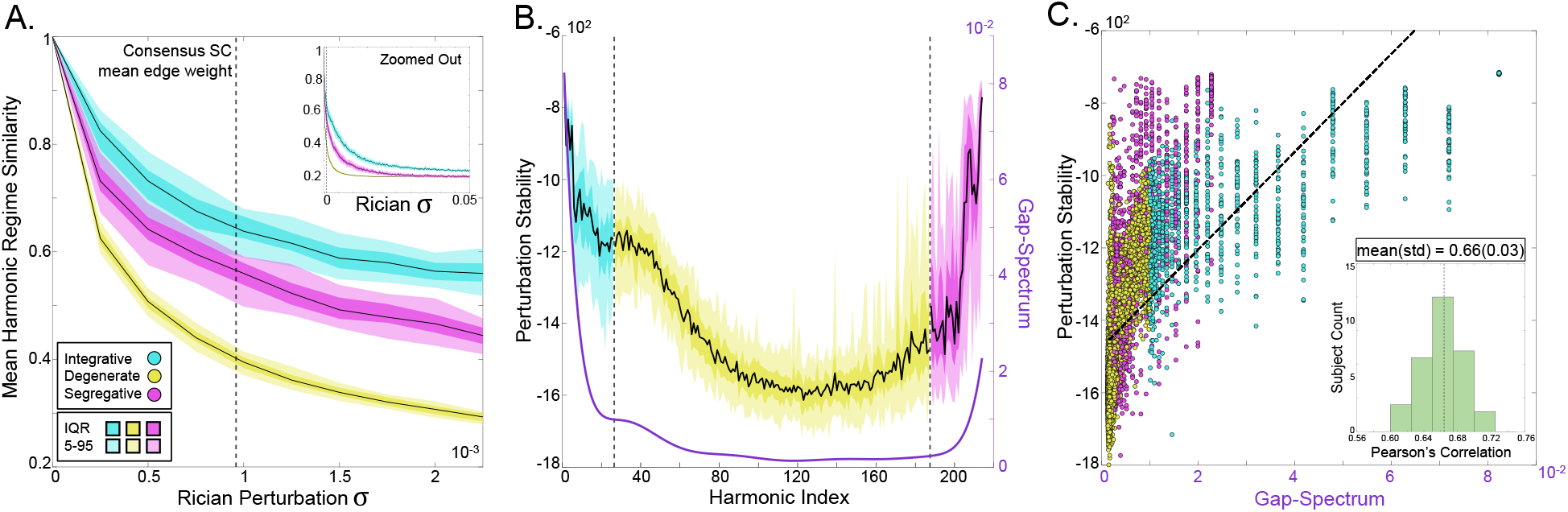
Stability of Harmonics. We perturbed the natural SC network across subjects with Rician noise over a wide range of *σ* (i.e., standard distribution). (A) We plot each regime-averaged intra-subject similarity between the authentic harmonics and the perturbed harmonics. The black line represents the median, while the darker and lighter shaded regions represent the IQR and 5-95 percentile range, respectively. Integrative harmonics show the most stability, segregative harmonics show intermediate stability, and degenerate harmonics show the least stability. The inset shows the same perturbation effect, but over a wider range of *σ*. (B) We quantify the stability of each harmonic and show that the stability follows the same general shape as the consensus gap-spectrum. (C) A scatter plot with data from all subjects and all harmonics, showing the overall relationship between the stability of harmonics and the gap-spectrum. The inset shows the distribution of subject-specific correlations between the stability of harmonics and gap-spectrum.

We then quantified the stability of harmonics (see Methods), and we found that stability indeed followed the same shape as the gap-spectrum (Figure 4B). The stability of harmonics across subjects was significantly correlated with the consensus gap-spectrum (mean *r* = 0.66; all *p <<* 0.001; Figure 4C).

### Regime-specific SC accounts for integrative and segregative graph theory metrics

We assessed whether regime-specific SC had graph theory metrics commonly related to integration and segregation. Qualitatively, the integrative SC shows broad connectivity across regions, the degenerate SC retains much of the connectivity structure from the native SC, and the segregative SC is sparse but dense in small clusters. Quantitatively, each regime SC embodies different graph metrics: 1-way ANOVAs showed significant regime-wise differences for all graph metrics (all *p <<* 0.01, Bonferroni corrected), and post-hoc two-tailed t-tests showed that most pair-wise differences were highly significant (*p <<* 0.01, Bonferroni corrected) (Figure 5). Modularity was greatest for the full network and equally large for degenerate and segregative regime SC, suggesting that modularity emerges from both degenerate and segregative harmonics. Global efficiency was greatest for the integrative SC, indicating that integrative harmonics facilitate efficient information transfer through the network. Small worldness was largest for the full SC, and second largest for the degenerate SC (the difference was small but still significant *p* = 7.3 *×* 10^−17^), suggesting that degenerate harmonics account for much of the small-worldness observed in the full network. Finally, coreness was greatest for the segregative SC, revealing that segregative SC is characterized by highly clustered modules.

**Fig 5:**
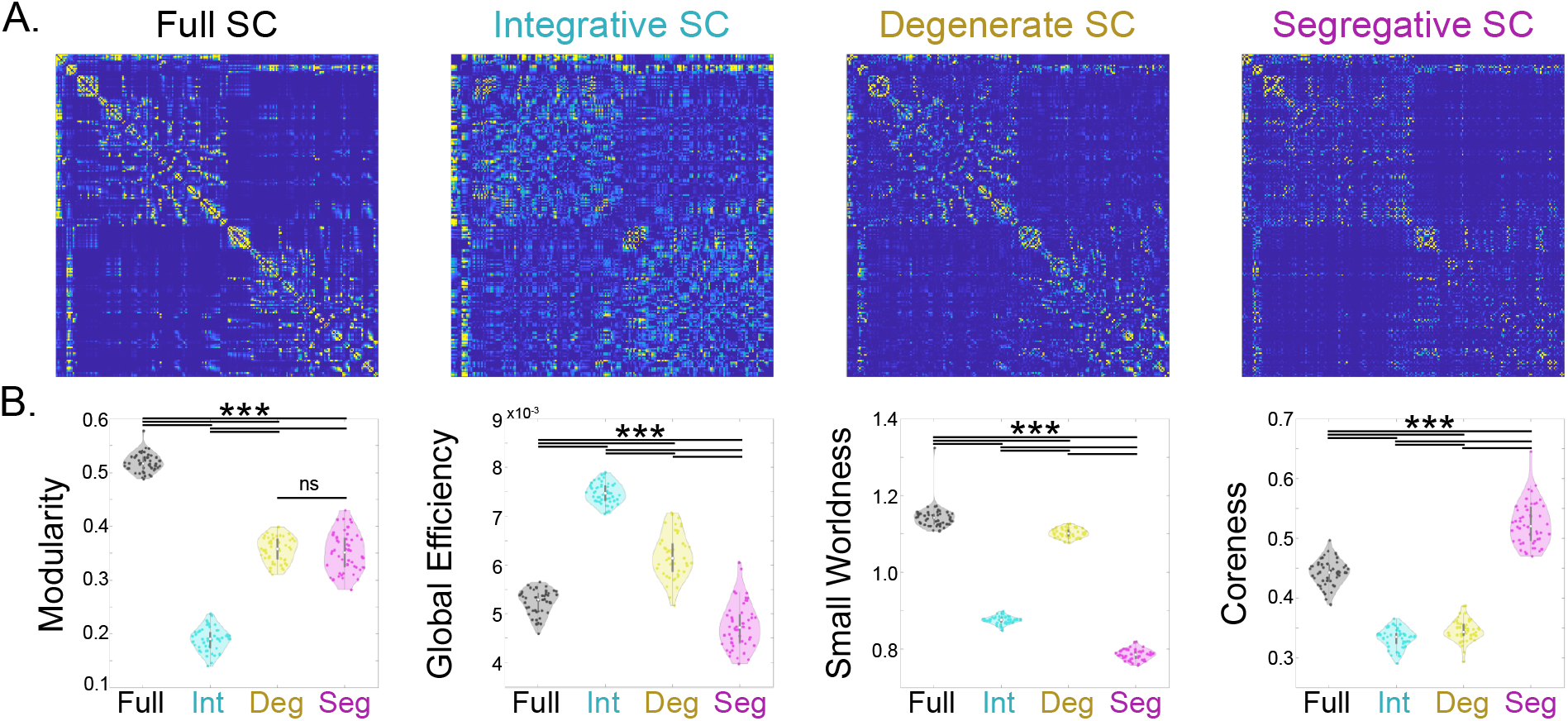
Regime-Specific SC. (A) We show (left to right) the consensus SC and its decomposition into integrative SC, degenerate SC, and segregative SC. (B) Violin plots of graph theory metrics computed for the full SC and each of the low-rank SC networks. From left to right: modularity, global efficiency, small worldness, and coreness. (*** indicates *p <<* 0.001, Bonferroni corrected; ns indicates non-significance).

### Harmonics have diverse participation with RSNs and functional tasks

We next assessed how SC harmonics related to functional resting-state networks (RSNs) and metaanalysis task activation maps. We found that RSNs show regime specificity across subjects (Figure 6A&C). The ternary plot showed that the Control Network and the Salience Ventral Attention Network participated most in the degenerate regime; the Subcortex, Visual, and Limbic Networks participated most in the segregative regime; the Somatomotor and Dorsal Attention Networks participated in a balance of the degenerate and segregative regimes; and the Default Mode Network was balanced between all three regimes (Figure 6C). RSN participation followed a similar relationship across Schaefer parcellation scales (Supplement Figure S9). We further found that whole brain task activation maps correlated with SC harmonics (Figure 6B). The ternary plot showed that tasks also had differential participation in all three harmonic regimes (Figure 6D). For example, Prediction and Learning was mostly integrative, while Inhibition was mostly degenerate, and Emotion was mostly segregative. However, most cognitive tasks involved a balance of the three regimes, such as Language and Decision Making, having approximately 50% integrative, and between 20-30% degenerate and segregative.

**Fig 6:**
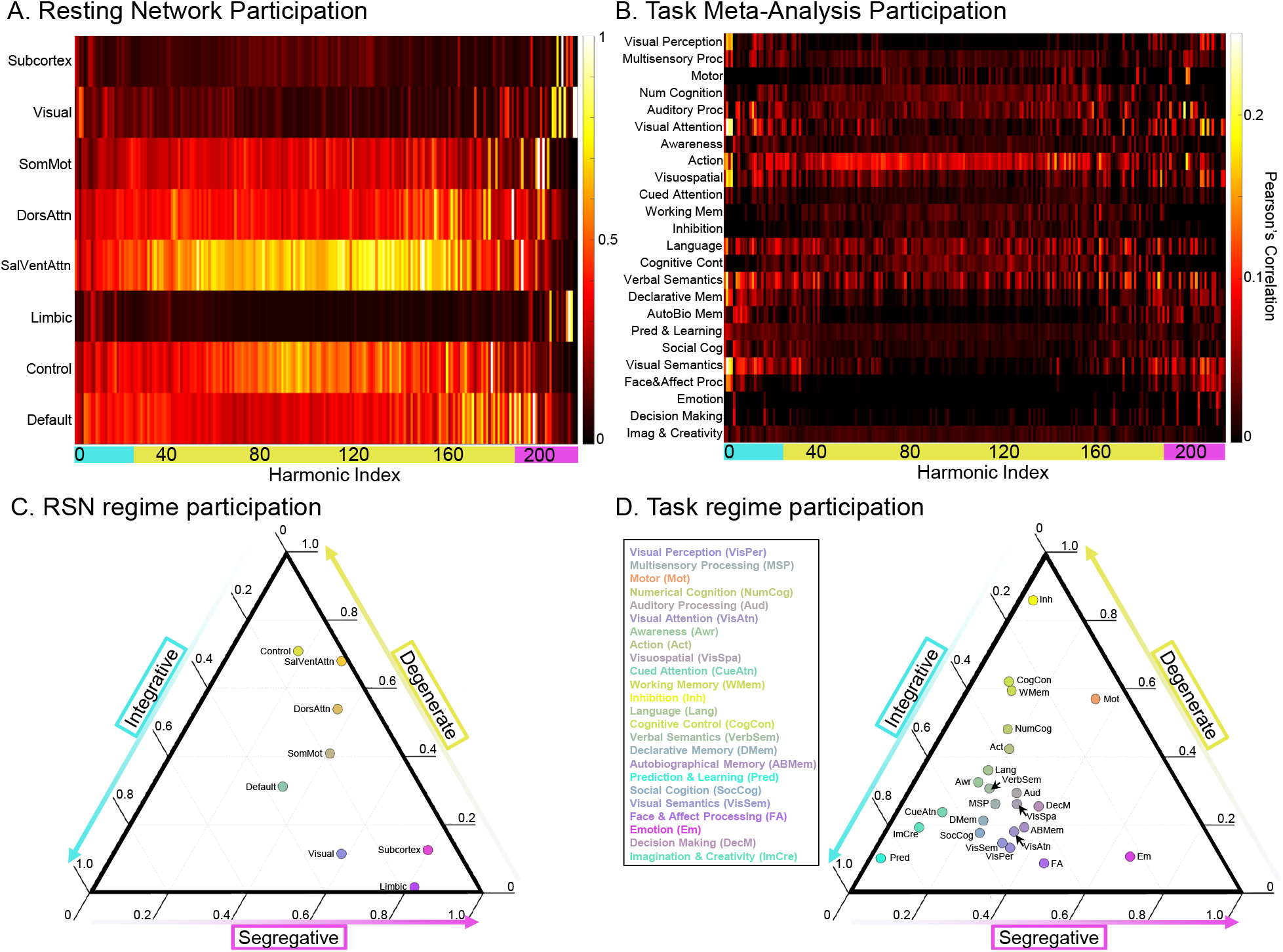
RSN and Neurosynth Coupling. We evaluated which harmonics had spatial organization that coupled with (A) resting-state functional networks (RSNs) and (B) Neurosynth task activation maps. All harmonics show spatial patterns that aligned with known functional organization. Below (C & D), we show ternary plots representing the above heat-maps as relative mixtures of each regime’s participation in (C) RSNs and (D) Neurosynth task activation networks. Note that the vertices of a ternary plot represent full participation in the respective regime, and the axis ticks are slanted to indicate the direction of the axis along dashed lines. Note that the DMN, for example, is nearly at the center of the RSN ternary plot, meaning it has close to equal participation from all three regimes. Conversely, the Limbic network showed almost exclusive participation in the segregative regime. The bubbles inside the ternary plot are colored based on their proximity to each of the three vertices (cyan–integrative; yellow–degenerate; magenta–segregative).

### Harmonic projection resting-state activity is different across regimes

Having found a relationship between harmonic regimes and known functional organizations, we hypothesized that empirical resting-state activity would also show similar regime-specific differences and relative regime contributions. We evaluated the functional characteristics of the three regimes in resting-state fMRI data projected onto the harmonic power vectors (Figure 7 A).

**Fig 7:**
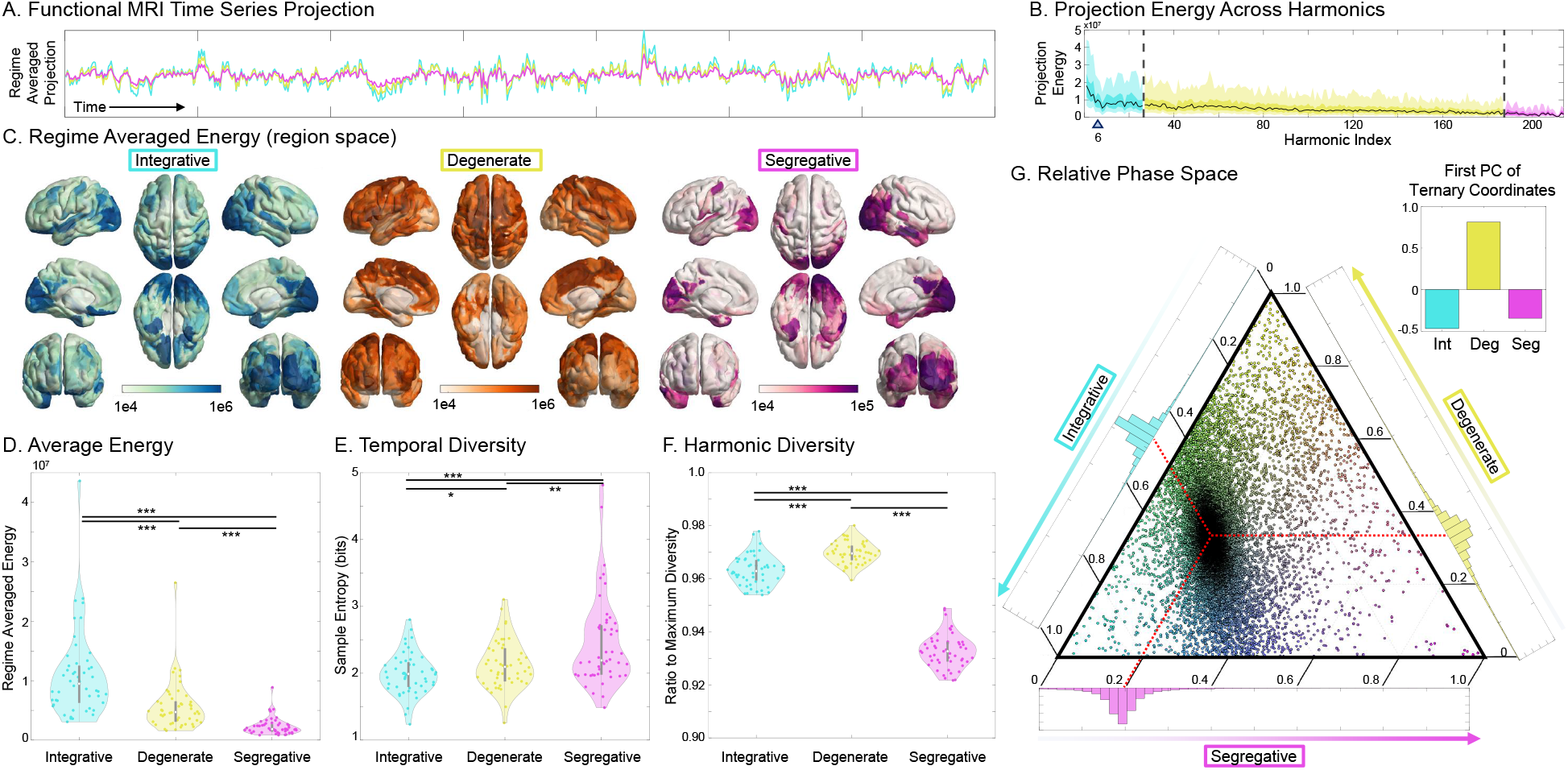
Regimes in Resting-State fMRI. (A) We project the the resting-state functional MRI time series onto the harmonic power vectors and show the regime-averaged projection for a single subject. (B) The projection energy (L2-norm across time) across all SC harmonics. The blue triangle indicates the previously reported divide between coupled/decoupled harmonics^[21]^. (C) The projection energy is mapped back to region space, showing region-wise resting-state functional participation in each regime. (D) Quantifying the regime-wise projection energy showed that the integrative harmonics have the most energy per harmonic, followed by the degenerate harmonics, and least for the segregative harmonics. (E) Quantifying the harmonic signal’s temporal diversity (i.e., sample entropy) showed the opposite pattern, with segregative harmonics carrying the most bits per harmonic, followed by degenerate and integrative regimes. (F) Quantifying the harmonic diversity of each regime in the functional time series showed that many degenerate harmonics are co-active, while integrative and segregative harmonics are more specific in time. (G) With a ternary plot, we show that the brain spends most of its time in a state of 46.5% integration, 33.6% degenerate, and 19.9% segregation. The upper-right inset shows the first principal component of this distribution, indicating that most functional variation is along the degenerate axis. (*:*p <* 0.05, **:*p <* 0.005, ***:*p <<* 0.001)

We quantified three key properties of the functional projection: projection energy, temporal diversity, and spatial diversity. The regime projection energy (L2-norm squared across time) gradually decreases from low to high harmonics with no clear demarcation between regimes (Figure 7 B). The blue triangle indicates the previously reported divide between coupled/decoupled harmonics based on the reported eigenvalue^[21]^; this showed that the previously identified “decoupled” regime is partially within our integrative regime and entirely within the degenerate and segregative regimes. Quantifying the spatial energy distribution across brain regions showed that integrative harmonics had a broad energy distribution across the cortex, with energy peaks located in sensory cortices; degenerate harmonics had energy weighted highest in frontopariteal and inferior temporal lobes; and segregative harmonics had sparse energy, with clusters of high energy in sensory regions, temporal poles, and ventromedial/orbitofrontal cortices (Figure 7 C). Overall, each regime had significantly different energies, with integrative having the most, segregative having the least, and degenerate in-between (all *p <<* 0.001; Figure 7 D).

We next analyzed temporal diversity (sample entropy) and spatial diversity (harmonic entropy) across regimes. The average temporal diversity contained in each harmonic regime projection showed that the segregative harmonics contained the most temporal diversity compared to either integrative or degenerate regimes (*p <* 0.004; Figure 7 E). Degenerate harmonics also had significantly more temporal diversity compared to integrative harmonics (*p* = 0.04). Regimes also showed significantly different harmonic (spatial) diversity, with the degenerate regime having the most diversity, followed by the integrative regime, and least in the segregative regime (all *p <<* 0.001; Figure 7 F).

Finally, we analyzed dynamics in the relative participation between regimes. We found that resting-state function across subjects had a strong tendency to be around approximately 46.5% integrative, 33.6% degenerate, and 19.9% segregative (Figure 7 G). With principal component analysis, we found that the degenerate regime formed the principle axis of temporal variation (Figure 7 G inset).

## Discussion

In this work, we have shown that an integration-segregation continuum in SC can be identified through its graph Laplacian harmonics and a feature we call the “gap-spectrum.” The gap-spectrum represents a fundamental feature of SC that naturally divides harmonics into three regimes: integrative, degenerate and segregative. This harmonic trichotomy reveals distinct regime-specific roles for inter-subject agreement, stability of harmonics, and functional relevance. As we discuss below, this proposed integration-segregation continuum subsumes, refines, and expands upon prior work on SC harmonics and integration/segregation in brain networks, therefore unifying these previously disparate areas of network neuroscience research.

Research on the brain’s graph Laplacian harmonics is a recent but actively growing area of computational network neuroscience^[12]^. Brain harmonics research encourages a conceptual transformation from thinking of the brain in terms of individual regions to spatially distributed patterns of structural connectivity. Indeed, this transformation has fundamental mathematical relationships to the Fourier transform^[15]^, and this motivates further borrowing intuitions from Fourier theory, such considering SC Laplacian eigenvalues as “spatial frequencies”^[12,24,54]^. These high eigenvalue harmonics represent bipartite-like subnetworks^[41,62,63]^, which could be interpreted as a form of “high frequency” network oscillation. While these intuitions are well-motivated, our results suggest that not all Fourier intuitions perfectly apply to brain networks. Notably, we show that higher “frequency” eigenvalues don’t necessarily correspond to higher spatial frequencies, but are instead smooth across large parts of the brain except for in small, localized clusters, suggesting they are “segregative” in nature. Fourier intuitions may also suppose that these higher harmonics are noisy, yet we have found that segregative harmonics are both preserved across subjects and stable to perturbation, with both of these properties showing a relationship with the gap-spectrum. Critically, we replicated our spatial frequency and inter-subject agreement results in a separate independent dataset of 220 subjects with more diverse age demographics, imaged with different scanning parameters, processed with a different connectivity pipeline, and parceled in a different atlas, all of which strongly suggests that our findings have strong generalizability. Together, this emphasizes the need for future work to carefully evaluate the extent to which intuitions from Fourier theory apply to brain harmonics.

Despite some departures from conventional Fourier wisdom, this “graph Fourier transform” still mathematically represents how signals would propagate though a network, therefore offering a promising way to bridge our understanding of the brain’s structure and function. In general, past work has implemented two different approaches with Laplacian harmonics to understand the brain’s function. One approach emphasizes only the lowest eigenvalue harmonics^[25,14,24,64]^. This can be motivated for a variety of reasons such as parsimony^[65]^, the amount of explained variance^[14,24]^, or as the outcome of a model^[25]^. However, our present work suggests that much can be gained from analyzing all harmonics, since the brain’s function across many representations– RSNs, tasks, and empirical resting-state activity–can be understood as a balance of integrative, degenerate, and segregative regimes. Interestingly, no functional networks were exclusively integrative, and the only functional network to show a near-perfect balance of the three regimes was the DMN. We speculate that this could be related to the DMN’s position at the top of the functional hierarchy^[14]^, thereby allowing the DMN to facilitate state transitions between regimes. RSNs and tasks generally emphasize certain harmonic regimes more than others–functions related to sensation, sociality, and memory were balanced between integrative and segregative harmonics, while degenerate harmonics were related to complex “top-down” cognition, such as attention, inhibition, and working memory.

A second approach divides all SC harmonics into two categories based on their participation in resting-state fMRI time series (i.e., the equal energy split criterion)^[21]^. This framework identifies the lowest frequency harmonics as “coupled” and all higher harmonics as “decoupled” from function^[21,12,66]^. The coupled and decoupled components of the functional signal are commonly synthesized into a single metric known as the Structural-Decoupling Index (SDI)^[21]^, which divides a given region’s decoupled participation by its coupled participation. SDI has proven useful, demonstrating relationships with the mechanics of cognition and behavior^[21,67–69]^ and could serve as a bio-marker of development^[70,71]^ and disease^[72,73]^. Our results affirm the success of the coupling versus decoupling work while additionally suggesting a further refinement into the integrative, degenerate, and segregative trichotomy. From this new perspective, the SDI would be a measure of how much of function across the brain is weighted into degenerate and segregative harmonics versus integrative harmonics, with the effects likely driven by the degenerate harmonics since they occupy a significant majority of the decoupled modes. Since the degenerate harmonics are also the most subject specific, SDI could be considered a metric of subject-uniqueness in function, since it is effectively normalizing by the integrative harmonics, which are most preserved across subjects. However, we emphasize that in our proposed framework all harmonics “couple” with function in diverse ways related to the harmonic regime. A limitation of the coupling/decoupling approach is that harmonic classification depends on their interaction with brain function rather than on an inherent feature of SC, meaning that their functional relevance is partially entangled with their classification. By defining harmonics based on the gap-spectrum inherent to SC, we can study properties of harmonics and their functional relevance independent of their classification.

Other research on the brain’s structure-function relationship seeks to explain how the structural network facilitates integration and segregation through the lens of graph metrics. The brain’s integration is interpreted as efficient signal transmission combined with signal convergence to highly connected regions, while segregation is interpreted in terms of the brain network dividing into tight-knit communities or modules with specific functions^[6,7]^. However, there are two limitations with this view of integration and segregation. First, graph metrics often have an embedded (though subtle) notion of time; network efficiency, for example, measures how easily a signal can move between nodes to traverse full the network (i.e., greater efficiency implies less transmission time). Therefore, graph metrics measuring signal transmission and convergence can only account for integration in time, not in space, yet spatial integration is thought to be necessary to resolve the binding problem^[74]^. Second, graph metrics for integration and segregation are defined separately, and therefore, they are incommensurable to a single integration-segregation spectrum–this incommensurability is precisely the crux of the binding problem. The present work suggests an alternative view of integration and segregation as represented through graph Laplacian harmonic spectrum. These harmonics describe signal distributions in space, which resolves the first limitation, and their continuum on the eigenspectrum resolves the second limitation. Our proposed framework reveals that integration and segregation in the human brain is similar across subjects, stable to network changes, and is related to unimodal sensory networks. Moreover, we show that harmonic regime SC recapitulates intuitions from graph metrics measuring integration and segregation. Therefore, SC harmonics appear to subsume insights from the past frameworks while overcoming their limitations.

Using SC harmonics to define an integration-segregation spectrum also provides a natural def-inition for what lies between integration and segregation: the degenerate harmonics. Degenerate harmonics deviate from integration and segregation in many important ways: they are subject specific, they are sensitive to network changes, they embody the network’s small-worldness, and they relate to top-down functional networks. Degenerate harmonics have a sense of “flexibility” since they represent symmetries in the network^[44]^, they are related to network synchonizability^[45]^, and they can form linear combinations to produce new degenerate harmonics. This ties nicely into our finding that resting-state time series shows the greatest harmonic diversity and temporal variability in the degenerate regime, meaning that multiple degenerate harmonics are “co-active” and rapidly fluctuate on and off. This finding also aligns with recent work suggesting that FC constructed from “decoupled” harmonics are highly variable in time and may encode information in transient brain states^[75]^. We speculate that these functional characteristics may correspond to creating flexible linear combinations that combinatorially encoded contextual and “top-down” information. Degenerate harmonics being related to subject-specificity and frontoparietal networks also aligns with recent research on “brain fingerprinting,” which has highlighted the frontal and parietal cortices as those most related to subject uniqueness^[76–86]^ and neural plasticity^[83]^. Most promisingly, Griffa et al.^[87]^ has shown that decoupled FC (representing mostly degenerate harmonics) also contains highly unique features within frontoparietal networks useful for brain fingerprinting. Degenerate harmonics being flexible in this way also makes them an ideal substrate for neural plasticity when considering the brain’s frugal metabolic^[88]^ and wiring cost^[89]^ economy. These results together raise exciting possibilities for future work to use degenerate harmonics to investigate individual uniqueness and neuroplasticity.

Putting everything together, we propose that SC harmonics provide a framework to study, and perhaps help resolve, the long-standing binding problem in the philosophy of mind. In essence, the binding problem seeks to explain how the brain can bind segregated sensory channels into an integrated whole^[90]^. It has been shown that sensory cortices are more segregated from the rest of the brain compared to other regions^[91,92]^, and yet some unification of integration and segregation must be necessary to bring all senses into a single coherent precept^[93]^. The fundamental puzzle in the binding-problem is to find a brain mechanism that can coordinate signal flow and represent information at the global level while still leveraging regional/neuronal specializations. Such a brain mechanism should be fundamental to the brain’s structure, shared across individuals, and directly implicate sensory processing–our present work provides strong evidence that integrative and segregative SC harmonics accommodate all of these properties. It is also expected that candidate binding mechanisms should be related to states of consciousness^[90]^. SC harmonics again fulfill this expectation, with recent work showing that SC harmonic repertoire diversity in brain function increases while on psychedelics and decreases while sedated with propofol^[54]^. However, this raises a question: how do integrative and segregative harmonics interact to share information and produce this binding effect? While this question deserves careful attention in future research, we speculate that segregative harmonics may “activate” to incoming sensory stimuli, and segregative harmonics’ spatial overlap with integrative harmonics allows the signal information to spread between regimes and broadcast the newly “bound” information globally. We believe future work studying harmonics through this framework could illuminate many mechanisms of macroscopic brain function.

Of course, this work must be viewed in light of known limitations for SC harmonics. Noise, low resolution, and artifacts are common in diffusion MRI. Tractography algorithms and their associated parameters can greatly affect quality of tracts, including turning angle, stopping criteria, etc. These can have significant effects on connectome quality. Additionally, our choice of edge weights and the associated impact on the networks degree distribution may contribute to our reported results. While many prior studies have generally identified similar harmonics as shown here^[61,12,40]^, the degenerate regime may be particularly affected by these choices. Another limitation is our use of the consensus SC which was constructed from a cohort of 50 healthy young adult subjects. Future work will benefit from constructing a consensus SC with larger healthy samples, and it is presently unclear whether our findings would extend to clinical or developmental populations. With respect to our Neurosynth meta-analysis, the Neurosynth topic terms presented here were averaged across the LDA-400 list where we selected the 146 terms related to cognitive (but not illness or methodological) terms. We chose these to maximize similarity to previously published work related to SC and functional network harmonics^[14,21]^. We believed that averaging correlations across many LDA terms and subjects would be more likely to eliminate findings rather than cause spurious results. Finally, we want to acknowledge an inherent tension between presenting SC harmonics as an integration-segregation continuum while also defining three discrete regimes of that continuum. The three regimes depends entirely on defining the criteria for the regimes’ cut-off. We show that defining regimes based on the gap-spectrum curvature provides one way to distinguish different properties of SC harmonics, but this need not be the only effective method. Future work using Laplacian harmonics to study integration and segregation in the brain should define these regimes to fit the context of the particular scientific question being asked.

Ultimately, unifying integration and segregation into a single continuum is crucial to understanding systems-level processing in the brain. We propose a framework for this unification based on the graph Laplacian harmonics, which we show can account for an integration-segregation continuum along with a previously undefined center we term the “degenerate” harmonics. The proposed trichotomy of graph harmonics reveals previously unreported structure-function relationships in a manner that is not possible within the previous integration/segregation nor coupled/decoupled paradigms. This finding provides key biological and functional interpretations for structural brain harmonics, bringing network neuroscience closer to understanding subject-specific differences and a framework to resolve the long-standing binding problem.

## Code Availability

The code for all presented results is made publicly available on GitHub: (https://github.com/RajLab-UCSF/IntDegSeg).

## Data Availability

The dataset for this work comes from a publicly available resource provided in Royer et al^[34]^–see their article for details. For ease of use and reproducibility, we have provided a limited subset of these data in the code repository.

## Competing Interests

The authors declare no competing interests.

## Author Contributions

BS: Conceptualization, Data curation, Formal analysis, Investigation, Methodology, Software, Validation, Vizualization, Writing - original draft, Writing - review & editing. SN: Conceptualization, Methodology, Supervision, Writing - review $ editing. AR: Conceptualization, Methodology, Project administration, Supervision, Writing - original draft, Writing - review & editing.

## Notes

https://github.com/Raj-Lab-UCSF/IntDegSeg

### Competing Interest Statement

The authors have declared no competing interest.

### Summary of Updates

Major revisions include an analysis of Low rank structural connectivity constructed for each harmonic regime and results from a replication analysis. Additionally, we substantially updated the Methods section to enhance the descriptions of our analysis. We significantly revised the Discussion to better summarize the key findings and incorporate the new analyses.

