## Supplemental Material for "Integrative, Segregative, and Degenerate Harmonics of the Structural Connectome"

### 1 Replication Dataset

Two of the principal findings in this work are: 1) that harmonics do not strictly correspond to spatial frequency and 2) that harmonics with low and high eigenvalues have high inter-subject agreement. To ensure that these main findings were replicable, we used another dataset of SC collected with 220 healthy subjects (ages $18-75$; mean(std) $= 39.1 \pm 16.8$; $119$ women), and tested whether this dataset also showed the same relationships with harmonic spatial frequency and inter-subject agreement. This replication dataset was previously used as the healthy control sample for a recent study analyzing differences in tinnitus [6].

Our replication dataset has important differences compared to the MICA-MICs dataset:

1. The age demographics for our replication dataset are more diverse.
2. The replication dataset consisted of two different MRIs (one 3T Siemens scanner at the University of Minnesota, one 3T GE scanner at UCSF). The replication dataset followed the Human Connectome Project (HCP) protocol for structural and diffusion MRI, which is detailed elsewhere [4]. The MICA-MICs dataset does not follow the HCP protocol.
3. The SC generation pipeline was developed in-house and has systematic differences to the micapipe pipeline. The SC pipeline used for our replication dataset is described in detail here [1]. While our in-house pipeline follows many of the same steps as micapipe, perhaps the most important difference is that our pipeline is fully volumetric without mapping to the cortical surface, while micapipe requires mapping to the cortex with Freesurfer [2] before constructing connectivity. Another potentially important difference is in the streamline generation, where micapipe uses MRtrix3 [7] while our streamline generation uses DiPy [3]. Note however that both datasets used probabilistic tractography with SIFT-2 filtering.
4. The brain parcellation to define regions for SC is the Brainnetome atlas [5], which has many differences compared to the Schaefer atlas used in the MICA-MICs dataset. The Brainnetome atlas has a total of 210 cortical regions and 36 subcortical regions, all defined volumetrically. Conversely, the micapipe pipeline Schaefer atlases have many resolution (although in the main text we focus on the 200-region parceled cortex), and it appends 14 subcortical regions defined separately from the atlas.

Despite these differences, our underlying conclusions remain unchanged: (1) SC Laplacian eigenvalues do not directly correspond to spatial frequency, with the highest eigenvalues indicating segregative harmonics (Figure S 1); and (2) harmonic matching shows that the integrative and segregative harmonics are well preserved across subjects while the degenerate harmonics are most subject-specific (Figure S 2).

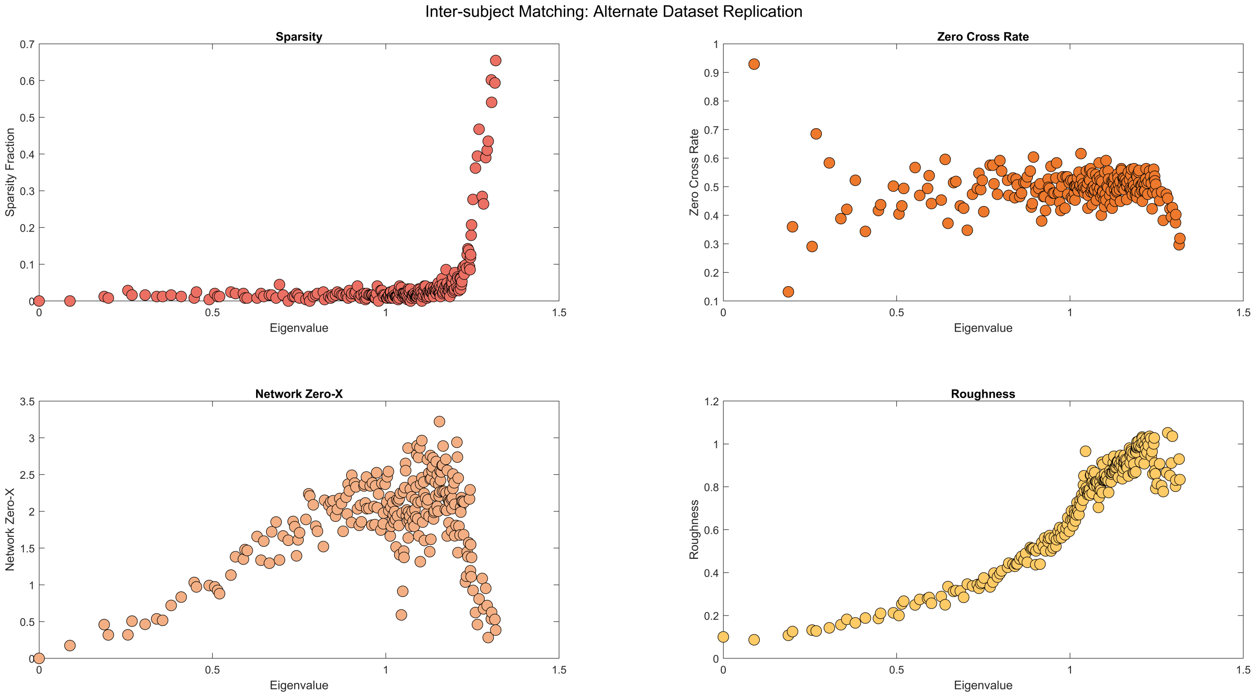


Figure 1: Harmonic frequency analysis of the SC consensus network generated as the mean across all 220 healthy subjects from the replication dataset.

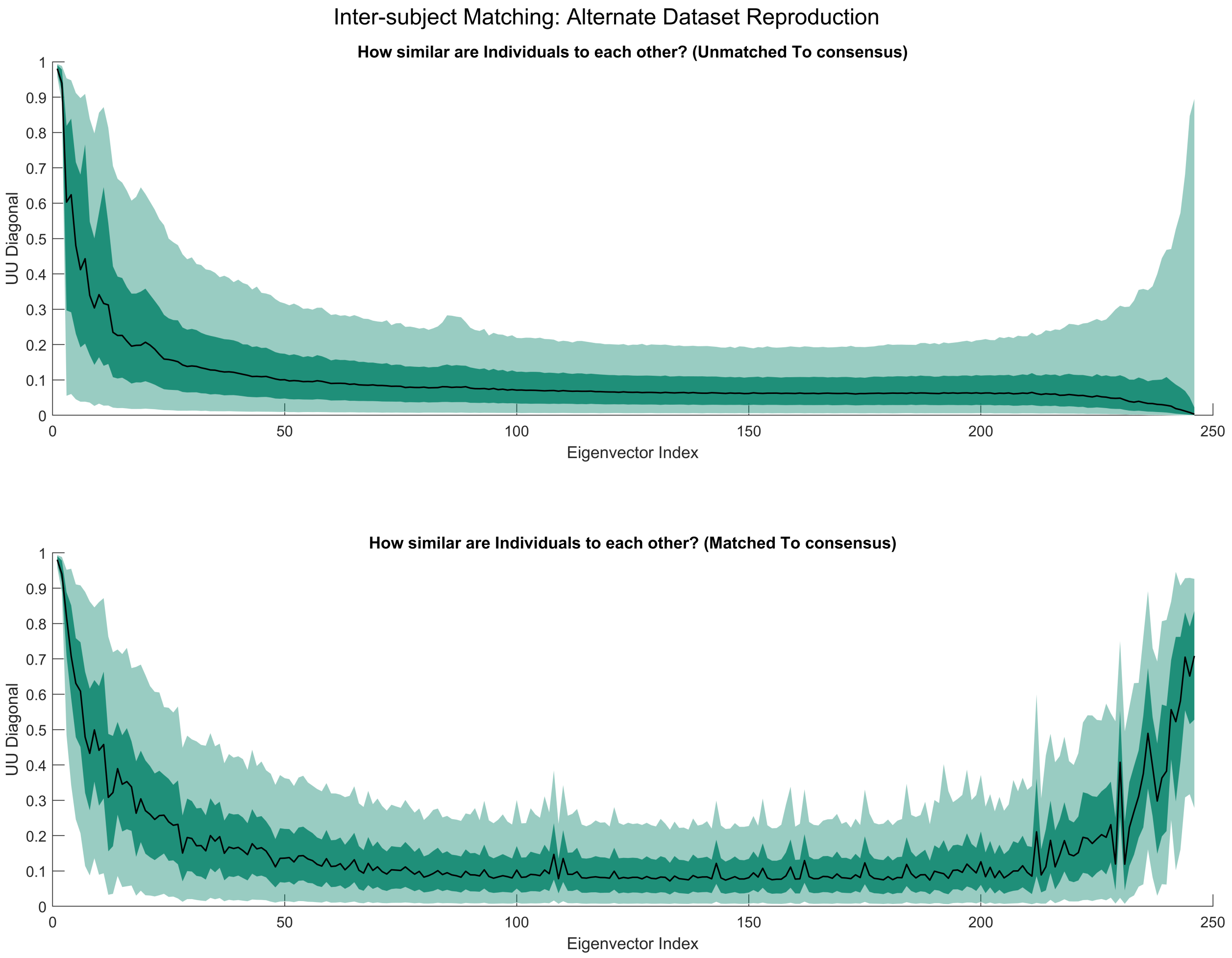


Figure 2: Inter-subject agreement analysis across all 220 healthy subjects from the replication dataset.

#### 2 Regional Adjacency Matrix

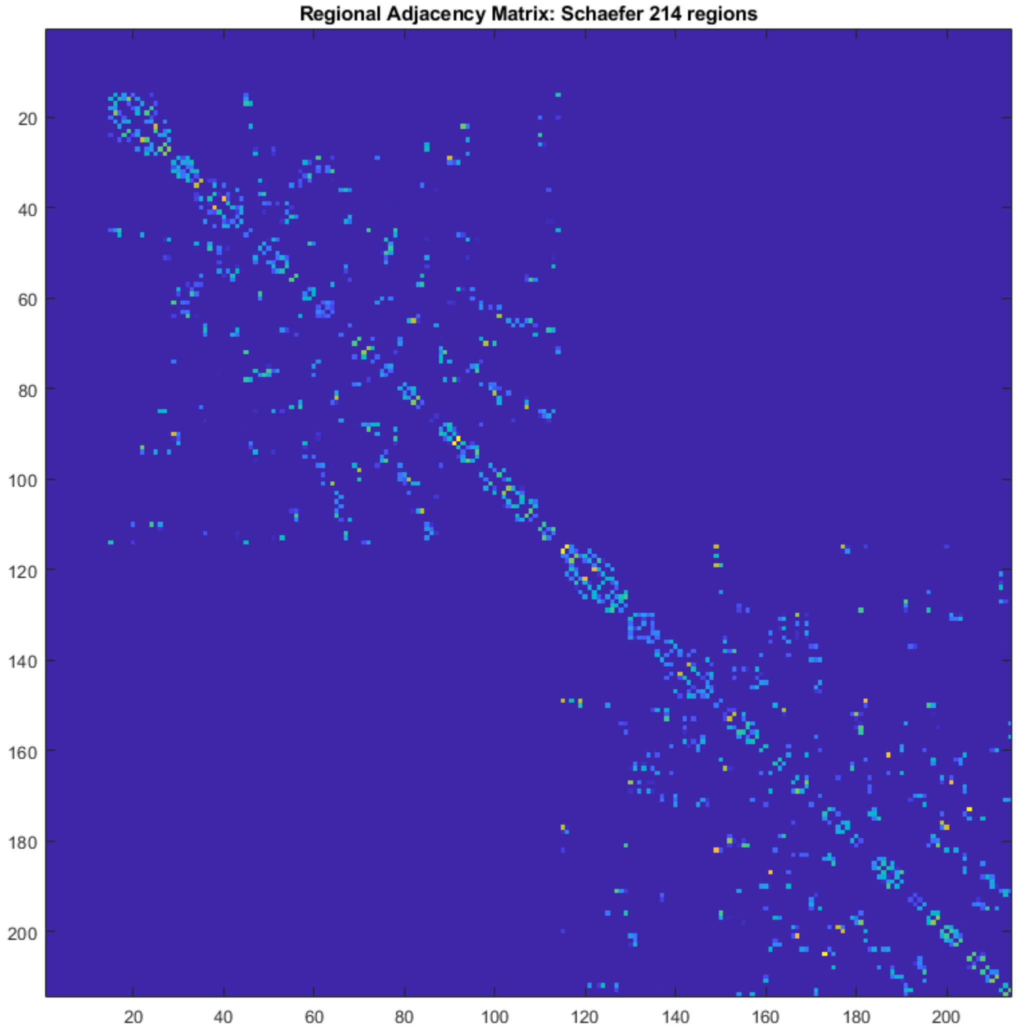


Figure 3: The Regional Adjacency matrix for the Schaefer 200 cortical parcellation. Entries of this matrix quantify the extent of adjacency between regions $i$ and $j$. Adjacency was computed by a summation of the inverse euclidean distance between adjacent voxels. Subcortex (comprising the first 14 rows/columns) is defined to have zero adjacency.

### 3 Network Zero-Crossings Across Thresholds

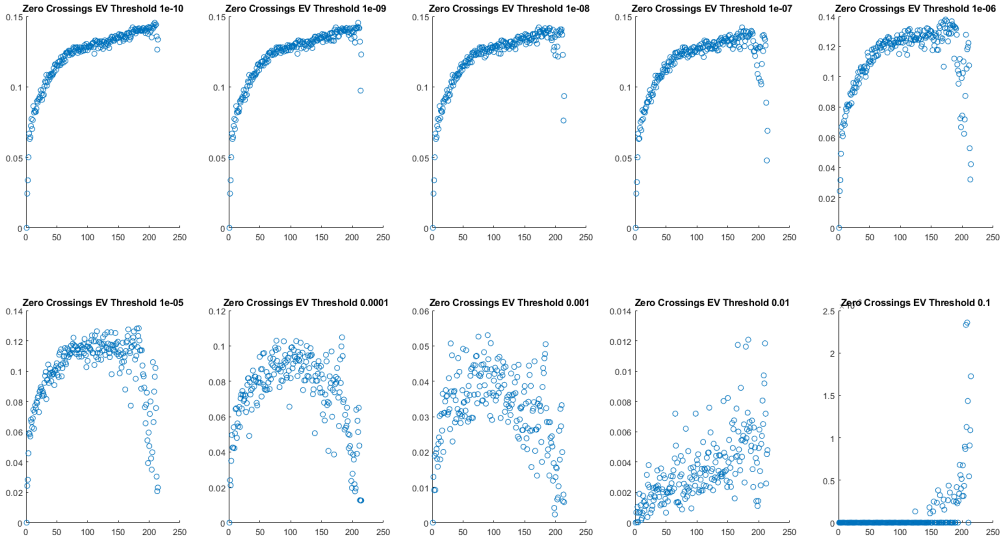


Figure 4: The Network Zero Crossings measure is sensitive to small fluctuations around zero (since it cares only for the sign, not magnitude, of the harmonic). For completeness, we show how this measure changes across various thresholds around zero.

### 4 Gap Spectrum Derivative for Regime Identification

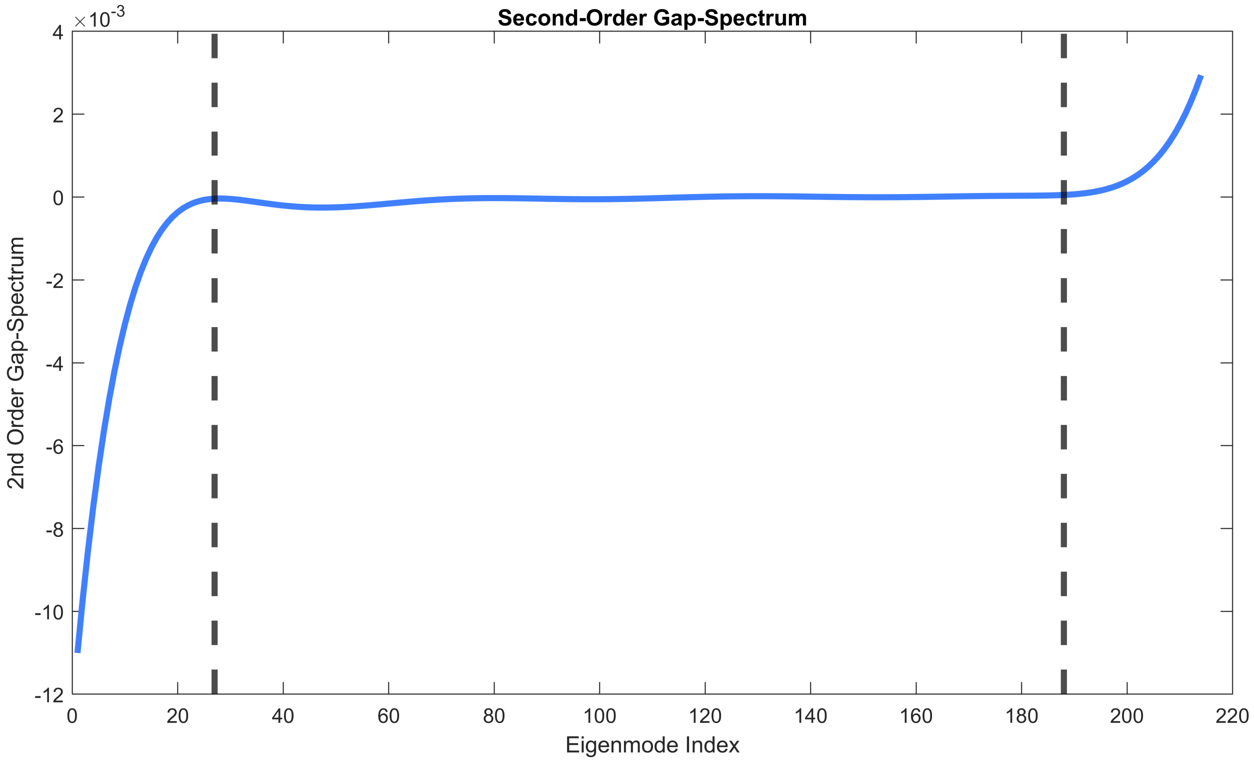


Figure 5: The Second-Order Gap-Spectrum, which was used to define the boundaries between the three regimes. Regime boundaries for each individual SC were identified as the first maxima (Integrative to Degenerate bound) and the last minima (Degenerate to Segregative bound). The final regime boundaries were based on the group-level boundaries after matching harmonics to the consensus SC, shown with black dashed lines.

### 5 Regime-specific (low-rank) SC Derivation

First, note that we have changed our notation from the main manuscript so that the standard Laplacian is $\mathcal{L}$ and the degree normalized Laplacian is $\check{\mathcal{L}}$, which will help clarify the following derivation.

Starting with the definition for the graph Laplacian of a network $C$ and a diagonal matrix of network degrees $D$:

$$\mathcal{L} = D - C \tag{1}$$

The low-rank truncation of any matrix by its eigenspectrum can be given by:

$$\tilde{\mathcal{L}} = \sum_{i=j}^{k} \lambda_i u_i u_i^T \tag{2}$$

Where the interval $[j, k]$ is a subset of harmonics. Suppose we truncate the Laplacian into $M < N$ subsets (where $N$ is the total number of network harmonics), then:

$$\mathcal{L} = \tilde{\mathcal{L}}_1 + \tilde{\mathcal{L}}_2 + ... + \tilde{\mathcal{L}}_M \tag{3}$$

We'll suppose that each $\tilde{\mathcal{L}}_i$ has a Laplacian form $\tilde{\mathcal{L}}_i = \tilde{D}_i - \tilde{C}_i$, therefore:

$$\tilde{\mathcal{L}}_1 + \tilde{\mathcal{L}}_2 + ... + \tilde{\mathcal{L}}_M = (\tilde{D}_1 - \tilde{C}_1) + (\tilde{D}_2 - \tilde{C}_2) + ... + (\tilde{D}_M - \tilde{C}_M) \tag{4}$$

And then note that

$$C = \tilde{C}_1 + \tilde{C}_2 + ... + \tilde{C}_M \tag{5}$$

Next, we perform the degree normalization of $\mathcal{L}$:

$$\check{\mathcal{L}} = D^{-1/2}\mathcal{L}D^{-1/2} = D^{-1/2}(\tilde{\mathcal{L}}_1 + \tilde{\mathcal{L}}_2 + ... + \tilde{\mathcal{L}}_M)D^{-1/2} \tag{6}$$

And therefore

$$\check{\tilde{\mathcal{L}}}_i = D^{-1/2}(\tilde{\mathcal{L}}_i)D^{-1/2} = D^{-1/2}(\tilde{D}_i - \tilde{C}_i)D^{-1/2} = D^{-1/2}\tilde{D}_iD^{-1/2} - D^{-1/2}\tilde{C}_iD^{-1/2} \tag{7}$$

Recall that when we compute the low-rank degree normalized Laplacian from its harmonics, we are computing $\check{\tilde{\mathcal{L}}}_i$, and then $\check{\tilde{D}}_i = \text{diag}(\check{\tilde{\mathcal{L}}}_i) = D^{-1/2}\tilde{D}_iD^{-1/2}$. To find $\tilde{C}$, we must rearrange and undo the degree normalization, giving us:

$$\tilde{C}_i = D^{1/2}(\check{\tilde{D}}_i - \check{\tilde{\mathcal{L}}}_i)D^{1/2} \tag{8}$$

Note that $\tilde{C}_i$ now can have negative off-diagonal entries where the original $C$ has strictly non-negative off-diagonal entries. These negative entries consist of network weights that must be canceled out in the full summation; therefore, they reflect a "negative image" of the network truncation. Our last step to recover the underlying graph representing the low-rank reconstruction is to remove all negative entries in $\tilde{C}_i$.

### 6 Resting State Network Supplemental Analysis

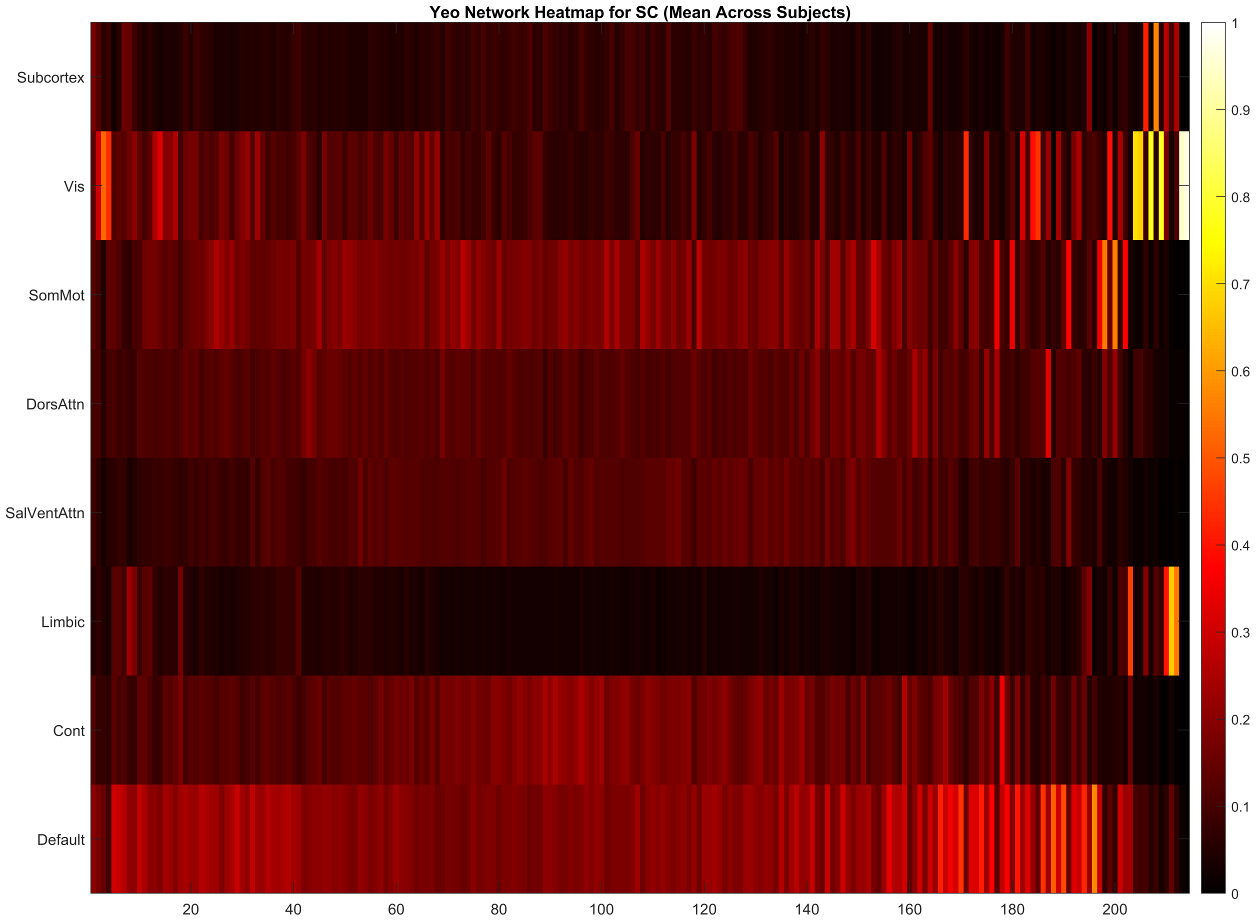


Figure 6: RSN participation without row-wise max-normalization.

### 7 Statistical Testing of Matching Performance

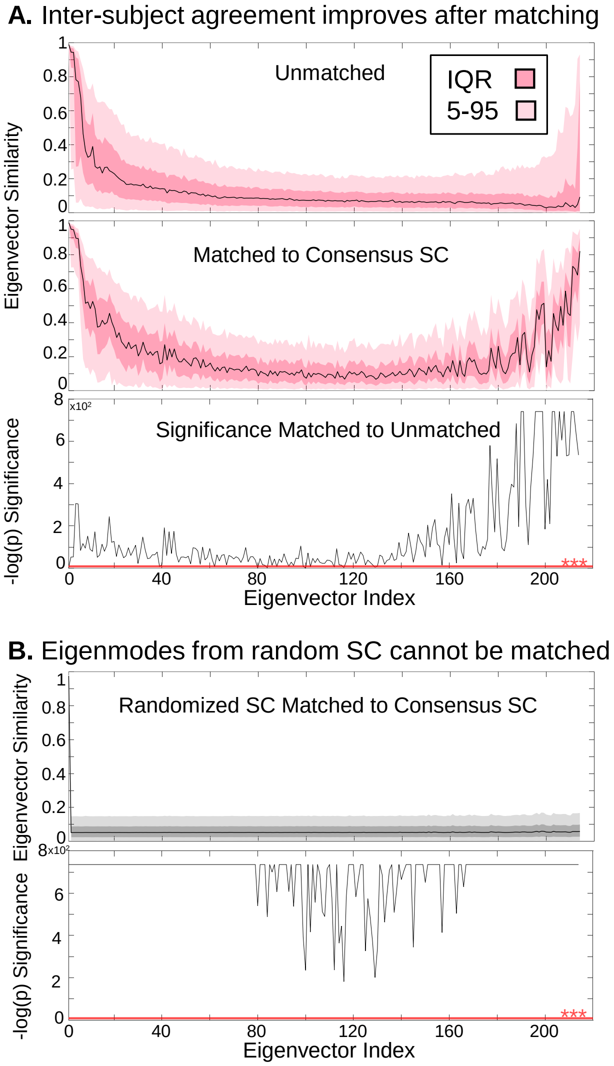


Figure 7: (A) Significance testing between the unmatched and matched distributions for each harmonic. The red line at the bottom of the Manhattan plot signifies the Bonferroni corrected significance threshold. All values above the red line indicate a highly significant difference between the unmatched and matched distributions. (B) Harmonics from a network with randomized edges matched to the original harmonics from the consensus SC network. The Manhattan plot beneath shows the significance between this randomized matching and the empirical matching. All subject-specific SC harmonics can match better to the consensus SC than harmonics from an edge-randomized network.

### 8 Temporal Frequency Supplemental Analysis

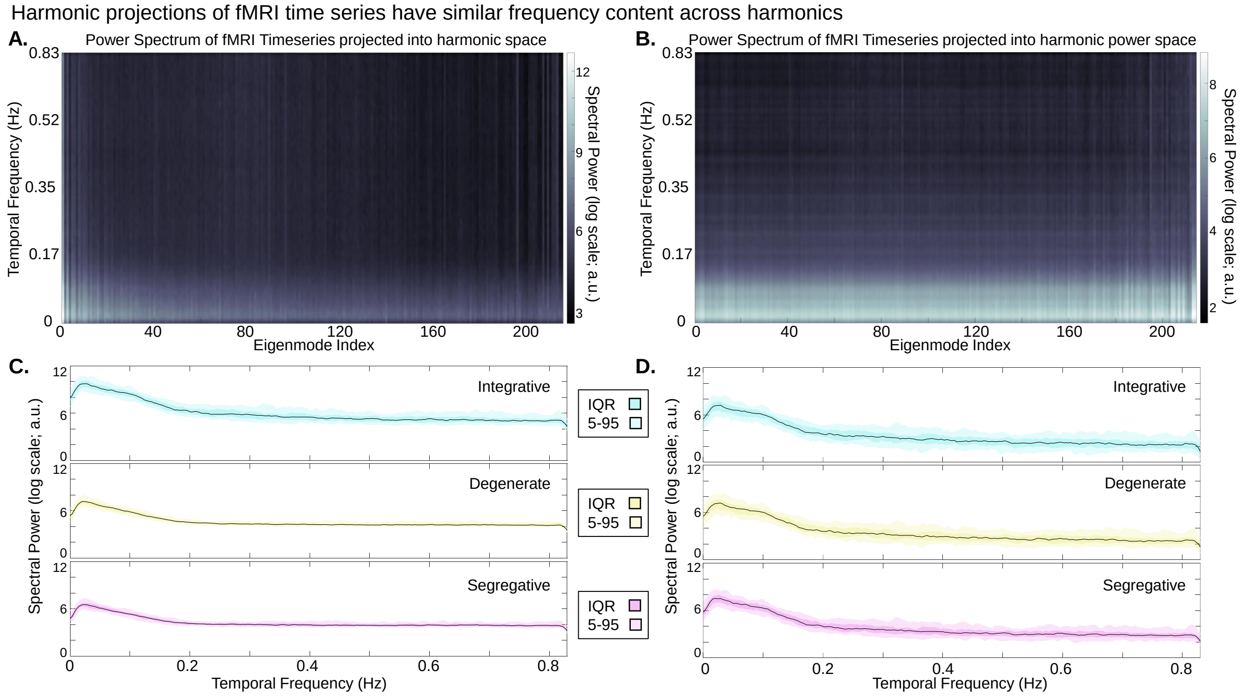


Figure 8: We compute measures for harmonic temporal frequency in relationship to the harmonics' participation in fMRI time series. We project the fMRI time series into the harmonic space (Graph Fourier Transform) and the harmonic power space (Equation 18 in main text). We use Welch's method to take the Fourier Transform of the resulting harmonic time series to analyze its frequency content. For completeness, we show both the spectral power for each harmonic projection using both the (A) bipolar as well as (B) power harmonic representations. (C & D) The temporal frequency characteristics are largely equivalent across all harmonics for both projections.

### 9 Inter-Subject Agreement Across Parcellation Scales

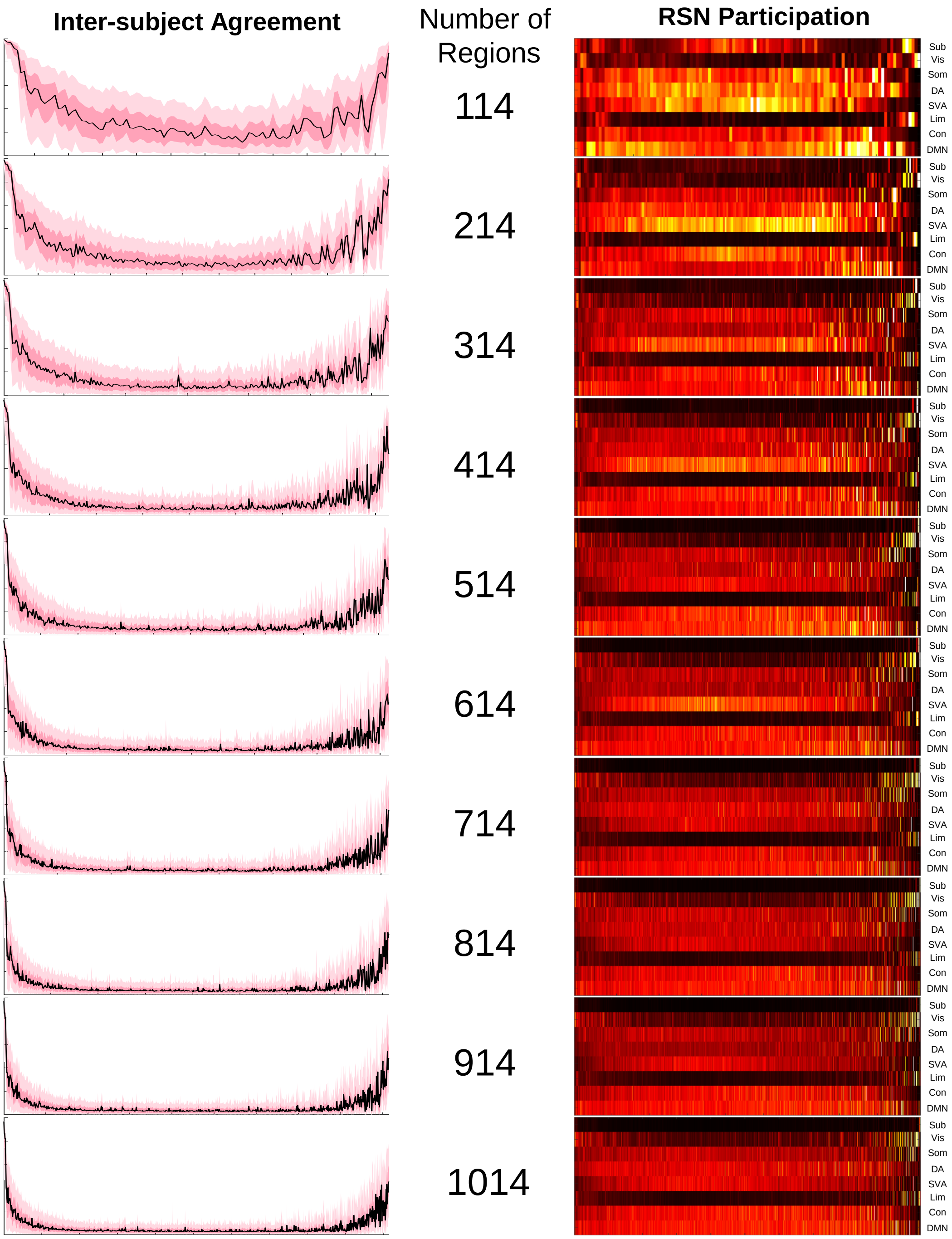


Figure 9: Inter-subject Agreement (left) and RSN participation (right) across all 10 Schaefer parcellation scales. General relationships at coarse parcellations qualitatively hold for fine parcellations.

### 10 Region-wise Inter-subject Variation

We examined which harmonics were most variable across subjects. We found that the segregative harmonics were the least variable between subjects compared to other harmonics ($p << 0.001$; Figure S10A). There was a general negative relationship between harmonic variation and mean inter-subject agreement ($r = -0.45; p < 0.001$; Figure S10A). We further found that the least harmonic variation across subjects were in the visual, somatomotor, auditory, and limbic areas (Figure S10B). Unexpectedly, the orbitofrontal cortex also showed low inter-subject harmonic variation. The areas with the greatest inter-subject variation were trans-modal functional network regions such as the bilateral parietal cortex and lateral prefrontal cortex (Figure S10B). Finally, we show the least variable Integrative and Segregative harmonics (Figure 10C).

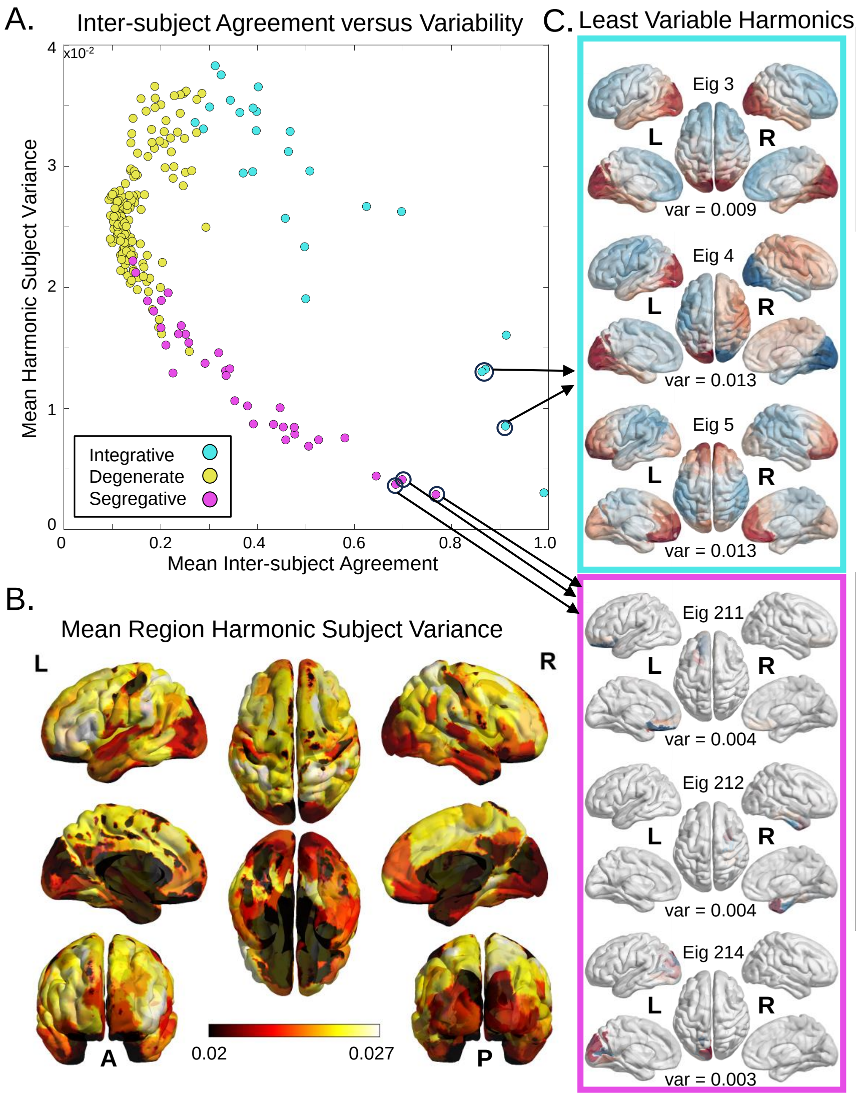


Figure 10: (A) The relationship between Inter-subject agreement and the variance of each harmonic across subjects. (B) Harmonic variance across subjects for each region. This shows that the sensory regions are the most similar across subjects in their harmonic representation. (C) A visualization of the least variable (i.e., most consistent) harmonics across subjects, selected from both the Integrative and Segregative regimes. (Note that the least-most variable Integrative harmonic is the trivial harmonic with eigenvalue = 0, which is not shown.)

### 11 Matching Permutation Distance

To further evaluate the harmonic matching, we computed the permutation length (defined as the number of column switches) for each harmonic across subjects, finding that the central harmonics required the longest permutations to match to the consensus ordering (Figure S11 top). We also computed the Kendall's Tau (KT) correlation, which is a measure of total permutation distance, for each subject with respect to the consensus SC (Figure S11 bottom), finding that overall subjects showed a matched harmonic ordering similar to the consensus SC (mean(std) KT = 0.90(0.015)).

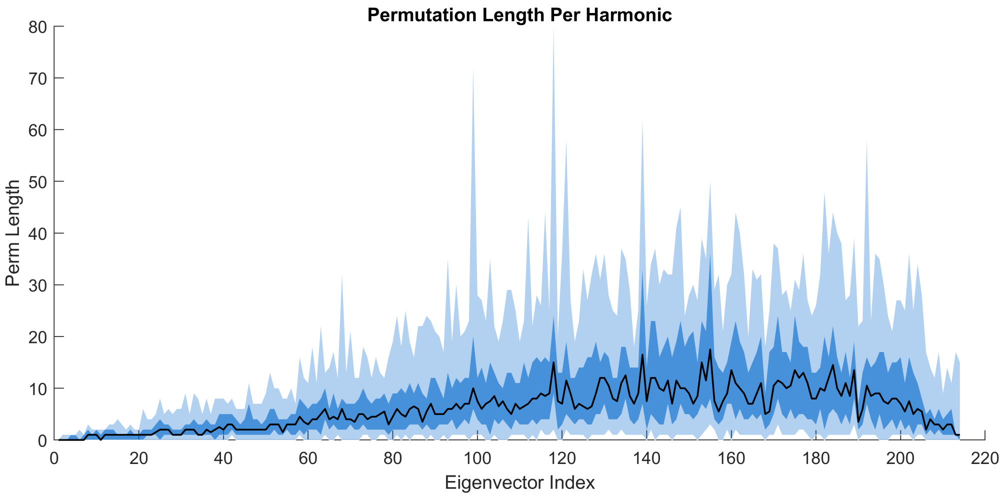


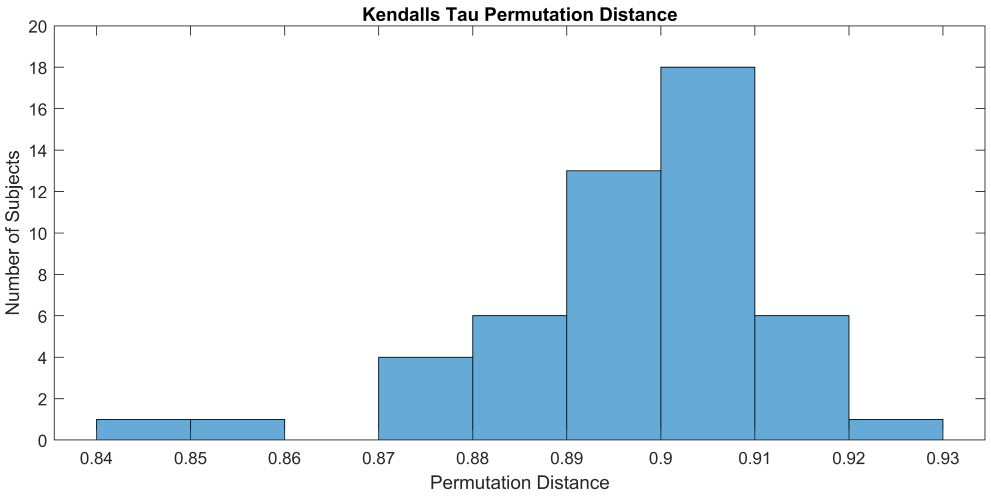


Figure 11: (Top) The permutation length (number of column switches) required to permute the harmonics. Darker regions denote the interquartile range (IQR) while lighter regions indicate the 5th to 95th percentile range. (Bottom) Kendall's Tau correlation between the permutation vector and an ordinal vector, which shows how far the permutation is from the consensus SC harmonic ordering.

### 12 Matching Alignment with Consensus

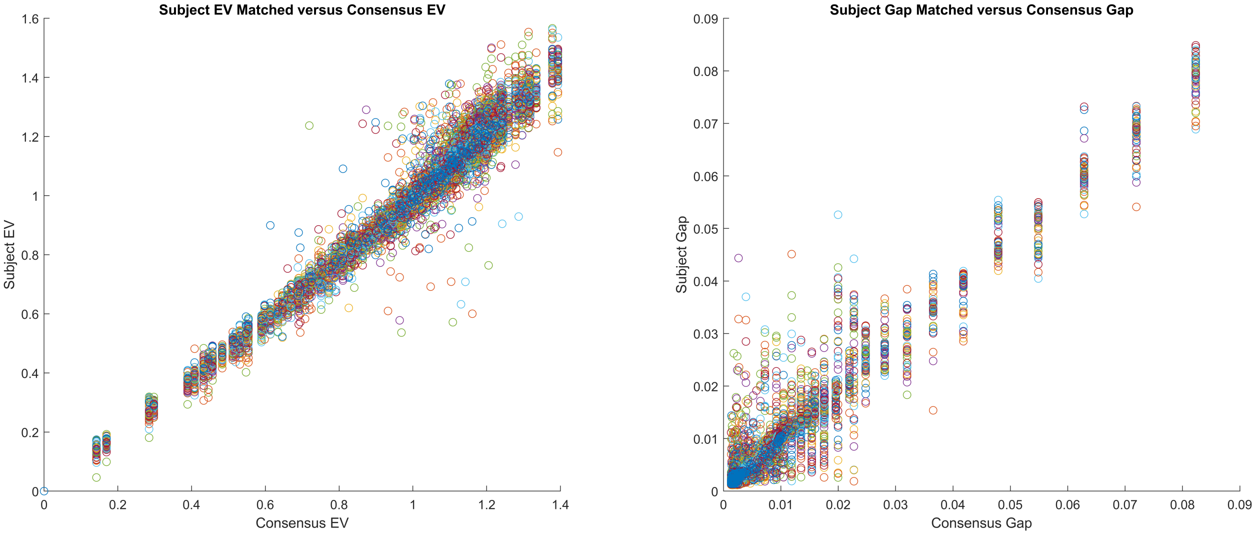


Figure 12: Subject-Specific eigenspectrum (left) and gap-spectrum (right) after matching compared to the consensus SC.
